# Target-enrichment sequencing reveals intergenotypic reassortment of infectious salmon anaemia virus in Atlantic Canada

**DOI:** 10.64898/2026.11.11.765878

**Authors:** Mingli Zhao, Jack O’Brien, Leighanne Hawkins, Ana Da Silva Filipe, Katherine Smollett, Silvia Soares, Michael Ness, Mari Aas Solheim, Nils Steine, Marius Karlsen, Sarah C. Hill

## Abstract

Infectious salmon anaemia virus (ISAV) causes severe disease and substantial economic losses in farmed Atlantic salmon (*Salmo salar*). The virus has an eight-segment, negative-sense single-stranded RNA (−ssRNA) genome and is divided into European and North American genotypes. Despite their co-circulation in locations such as the Atlantic coast of Canada, reassortment between these genotypes has never been observed. However, routine diagnostic and sequence-based surveillance often target individual genome segments and may therefore fail to identify reassortant viruses or accurately resolve their genomic ancestry. In this study, we developed a probe panel targeting both genotypes and applied Oxford Nanopore sequencing to 37 ISAV-positive samples collected from 14 farm sites in the Bay of Fundy, New Brunswick, Canada. Twenty-seven samples yielded 29 consensus genomes, including separate viral consensuses recovered from two mixed infections. Segment-specific phylogenies identified 24 viruses with a previously unreported intergenotypic genomic constellation comprising a European-genotype segment 3 and North American-genotype segments 1, 2 and 4–8. Their consistent monophyletic clustering across segment-specific phylogenies supports a single ancestral reassortment event followed by onward circulation of the reassortant lineage. A time-scaled phylogeny of segment 3 estimated the posterior mean time to the most recent common ancestor (tMRCA) of the sampled reassortant cluster as September 2016 (95% HPD: March 2013–June 2019), indicating that this segment 3 lineage pre-dated its first sampling in 2021. Five additional viruses represented several distinct EU-genotype lineages. To our knowledge, this is the first documented reassortment between the European and North American ISAV genotypes. These findings identify an intergenotypic reassortant lineage detected across multiple farms in the Bay of Fundy and demonstrate the value of whole-genome sequencing for detecting reassortment, mixed infections and epidemiologically relevant viral diversity.

## 1. Introduction

Reassortment is a major evolutionary mechanism in segmented RNA viruses, enabling the exchange of genome segments between coinfecting viral strains. It can generate novel viral genotypes with altered fitness, host range, pathogenicity or transmission potential (McDonald et al., 2016; Lowen, 2018). In aquaculture, reassortment is increasingly recognised as an important driver of viral emergence because high host densities, intensive farming practices and the global movement of fish and fish products facilitate the co-circulation and interaction of genetically distinct viral lineages (Valero and Cuesta, 2023). Reassortment has been reported in several segmented viruses that cause diseases of major economic importance in aquaculture, including infectious salmon anaemia virus (ISAV) (Devold et al., 2006; Cottet et al., 2010; Plarre et al., 2012). Yet the frequency and evolutionary consequences of reassortment in ISAV remain poorly understood, particularly whether divergent genotypes can generate viable reassortants capable of persisting and spreading in salmon populations.

ISAV belongs to the genus *Isavirus* within the family *Orthomyxoviridae* and has a segmented, negative-sense, single-stranded RNA genome (Mjaaland et al., 1997; Krossøy et al., 1999). Its genome comprises eight segments encoding structural and non-structural proteins, including the RNA-dependent RNA polymerase complex encoded by segments 1, 2 and 4; the nucleoprotein encoded by segment 3; the fusion (F) and haemagglutinin–esterase (HE) surface glycoproteins encoded by segments 5 and 6, respectively; non-structural proteins encoded by segment 7; and the M1 and S8ORF2 proteins encoded by segment 8 (Cárdenas et al., 2022). This segmented genome structure creates the potential for reassortment during coinfection, whereby progeny viruses may acquire genome segments from different parental strains. ISAV virulence is strongly associated with deletions in the highly polymorphic region (HPR) of the haemagglutinin–esterase (HE) gene. HPR0 variants retain the full-length HPR and are generally associated with transient, subclinical infections, whereas HPR-deleted (HPRdel) variants are associated with clinical infectious salmon anaemia (ISA) outbreaks in farmed Atlantic salmon (Mjaaland et al., 2002; Fourrier et al., 2014; Christiansen et al., 2017). ISA is one of the most consequential viral diseases affecting farmed Atlantic salmon and can result in high mortality, compulsory depopulation, disrupted production and substantial socioeconomic losses (Falk et al., 1997; Rimstad and Markussen, 2020).

ISAV isolates are generally divided into two major phylogenetic groups, the European and North American genotypes (Kibenge et al., 2007a; Madhun et al., 2019. European-genotype ISAV strains have caused large-scale epidemics in Europe, Chile and parts of the Americas, whereas North American genotype ISAV strains have historically been more geographically restricted but remain epidemiologically important in North America (Castro-Nallar et al., 2011; Kibenge et al., 2009; Kibenge et al., 2016; Madhun et al., 2019). Nevertheless, aquaculture practices, movements of fish and marine connectivity may facilitate the geographical spread and co-circulation of viruses from the two genotype groups (Valero and Cuesta, 2023).

In Canada, ISAV was first detected in 1996 in the Bay of Fundy and is now established along the Atlantic coast (Mullins et al., 1998). Major outbreaks between 1997 and 2007 caused substantial production and financial losses (Johnson et al., 2008). From 2012 onward, ISAV was again detected in Atlantic Canada, including in New Brunswick, Nova Scotia and Newfoundland and Labrador (Figure 1; Canadian Food Inspection Agency, 2026; Gagné and LeBlanc, 2018; Romero et al., 2022). Phylogenetic analyses indicate the co-circulation of endemic North American genotype viruses and introduced European genotype viruses (Gagné and LeBlanc, 2018). This co-circulation is particularly evident in Newfoundland and Labrador, where viruses of both genotypes have been detected at the same fish farm (Gagné and LeBlanc, 2018; Gautam et al., 2023). Understanding the recent resurgence of ISAV-HPRdel in Canada is important to protect both farmed and wild populations there, but few studies have investigated viral genetic factors that may be contributing to this.

**Figure 1.**
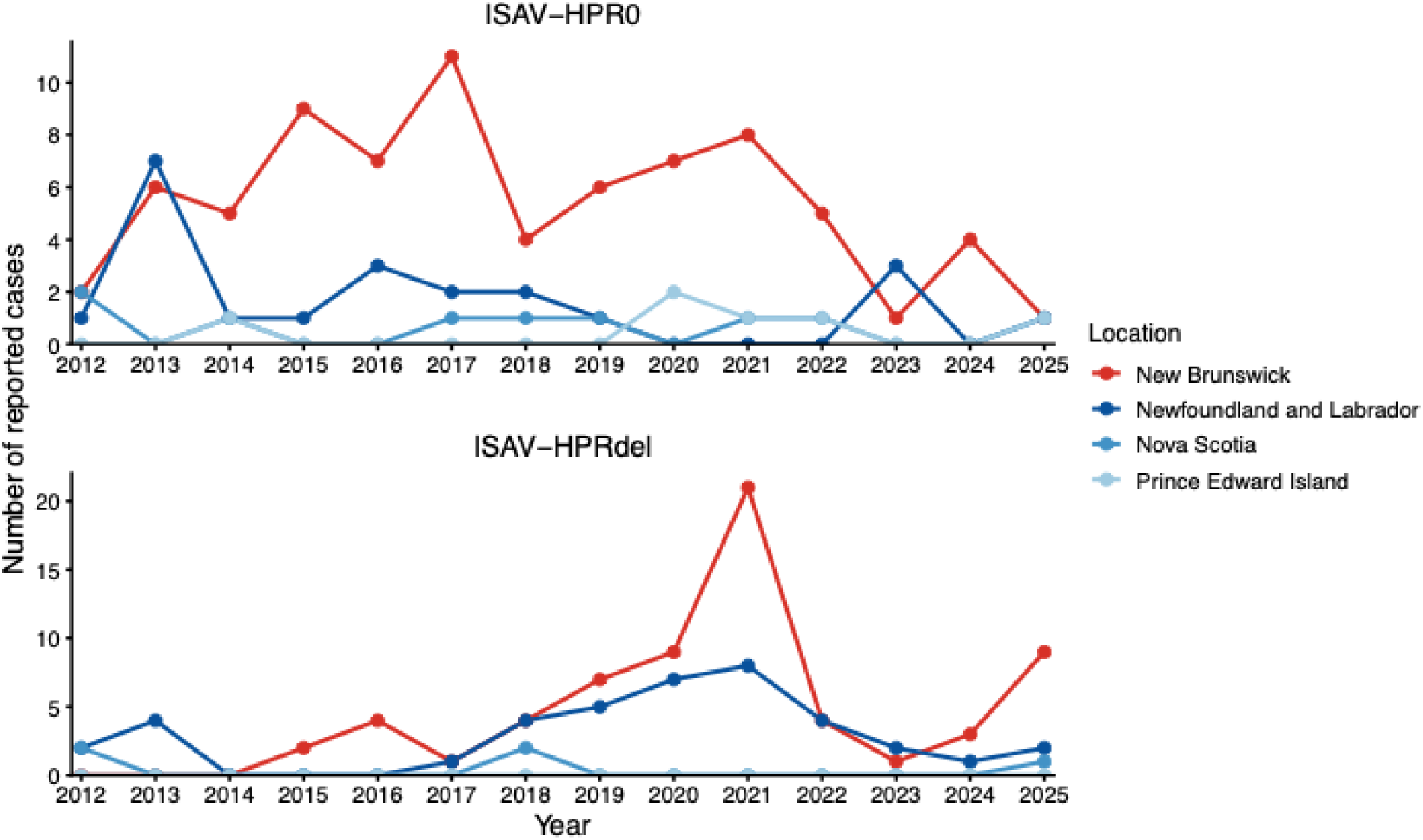
Temporal distribution of reported ISAV detections in Atlantic Canada by location, 2012–2025. Annual numbers of Canadian Food Inspection Agency (CFIA) ISAV records are shown for ISAV-HPR0 (upper panel) and ISAV-HPRdel (lower panel), with coloured lines indicating the reported location: New Brunswick, Newfoundland and Labrador, Nova Scotia, and Prince Edward Island. Records associated with the CFIA note “This virus strain is not known to cause disease” were classified as ISAV-HPR0, whereas all other records were classified as ISAV-HPRdel. Data from 2026 were excluded because the year was incomplete at the time of analysis (Canadian Food Inspection Agency, 2026).

Reassortment has likely contributed to the emergence and diversification of virulent ISAV lineages. For instance, circulating viruses during the Chilean ISA epidemic were identified as genomic mosaics derived from multiple European genotype ancestors (Cottet et al., 2010). In Atlantic Canada, co-circulation of genetically diverse European and North American genotype viruses creates opportunities for intergenotypic reassortment, but to date no convincing evidence of such reassortment has been identified. Publicly available ISAV sequence data remain highly uneven across genome segments and are dominated by European-genotype sequences, with relatively few isolates represented across all eight segments, limiting opportunities for genome-wide analyses of reassortment (Figure 2). Routine ISAV diagnostic and sequencing approaches often target segment 8 due to its high degree of genetic conservation, while further differentiation of HPR0 and HPRdel variants relies on segment 6. Sequencing of antigenic proteins encoded by segments 5 and 6 is common, yet partial sequencing of genomes is insufficient to detect discordant segment ancestries across the genome and hence reassortment (Munir and Kibenge, 2004). Although multiplex PCR can enable whole-genome sequencing of ISAV, the assay developed by Spilsberg et al. (2024) was designed for the European genotype and was not evaluated on North American viruses. This may limit detection of mixed-genotype infections and obscure evidence of reassortment.

**Figure 2.**
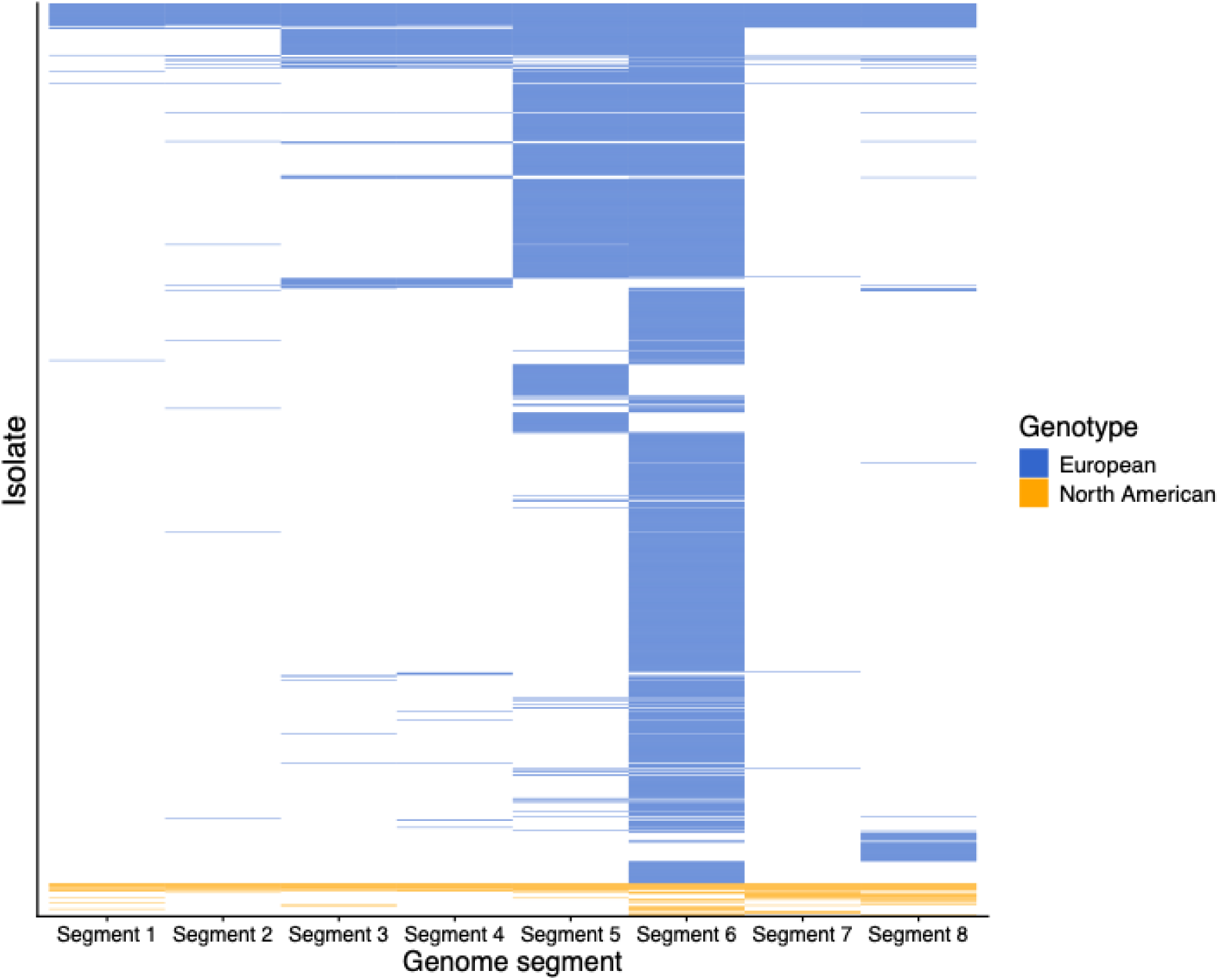
Availability and genotype classification of ISAV genome segments in GenBank (accessed 29 October 2025). Each row represents an ISAV isolate, and each column represents one of the eight genome segments. Coloured lines indicate the availability of sequence data for a given segment, but do not represent the length or completeness of the sequence. Colour indicates genotype classification, with blue representing the European genotype and orange representing the North American genotype. Isolates are sorted by European and North American genotype and, within each genotype, by the number of genome segments for which sequence data were available, in descending order.

To address these limitations, we developed a probe-capture hybridisation panel for whole-genome ISAV sequencing. Unlike fixed primer pairs, tiled oligonucleotide probes can enrich genetically divergent targets across broad sequence variation (Depledge et al., 2011; Weitemier et al., 2014). We applied this panel to ISAV-positive samples from the Bay of Fundy, New Brunswick to (i) assess genome recovery across a range of diagnostic Ct values; (ii) determine segment-specific genotype ancestry; (iii) detect mixed infections and reassortant genome constellations; and (iv) characterise amino-acid variation in the HE and F proteins.

## 2. Materials and methods

### 2.1 Study ethics

All Canadian samples used in this study were residual diagnostic specimens originally submitted to diagnostic laboratories through the New Brunswick ISAV surveillance programme. Under this programme, moribund fish were sampled across farm sites, with diagnostic testing conducted monthly when ISAV was not detected and every two weeks following detection of ISAV. Residual samples and associated metadata from commercial diagnostic laboratories were used with the written informed consent of the producers. Two ISAV-positive samples (North American genotype) provided by a Scottish governmental laboratory (Marine Directorate) were included as positive controls. Written informed consent from producers was not obtained for these controls; however, no identifiable producer-or site-level information was shared. The study was approved by the Royal Veterinary College Ethical Review Board (URN 2022 2098-3).

### 2.2 Samples and metadata

A total of 37 ISAV-positive samples from Canada were included. The samples comprised kidney tissue collected from adult farmed Atlantic salmon following disease outbreaks during the seawater production phase. Samples originated from 14 farm sites across two clearly separated production areas, designated here as A and B, in the Bay of Fundy, New Brunswick, and were collected between 2021 and 2024. The study area contains a high density of salmon farms, with each sampled farm located within 2 km of at least one other sampled or unsampled farm, and the maximum distance between sampled farms being less than 50 km. Between one and eight samples were obtained from each farm, with one to four samples collected from individual cages.

Diagnostic RNA extraction and reverse-transcription quantitative PCR (RT-qPCR) were performed by Pharmaq Analytiq AS, Norway using proprietary in-house techniques. Cycle threshold (Ct) values ranged from 13 to 37, representing a broad range of viral loads. Anonymised metadata for all sequenced samples are provided in Table 1; farm sites and production area information are anonymised to maintain anonymity of producers.

**Table 1.**
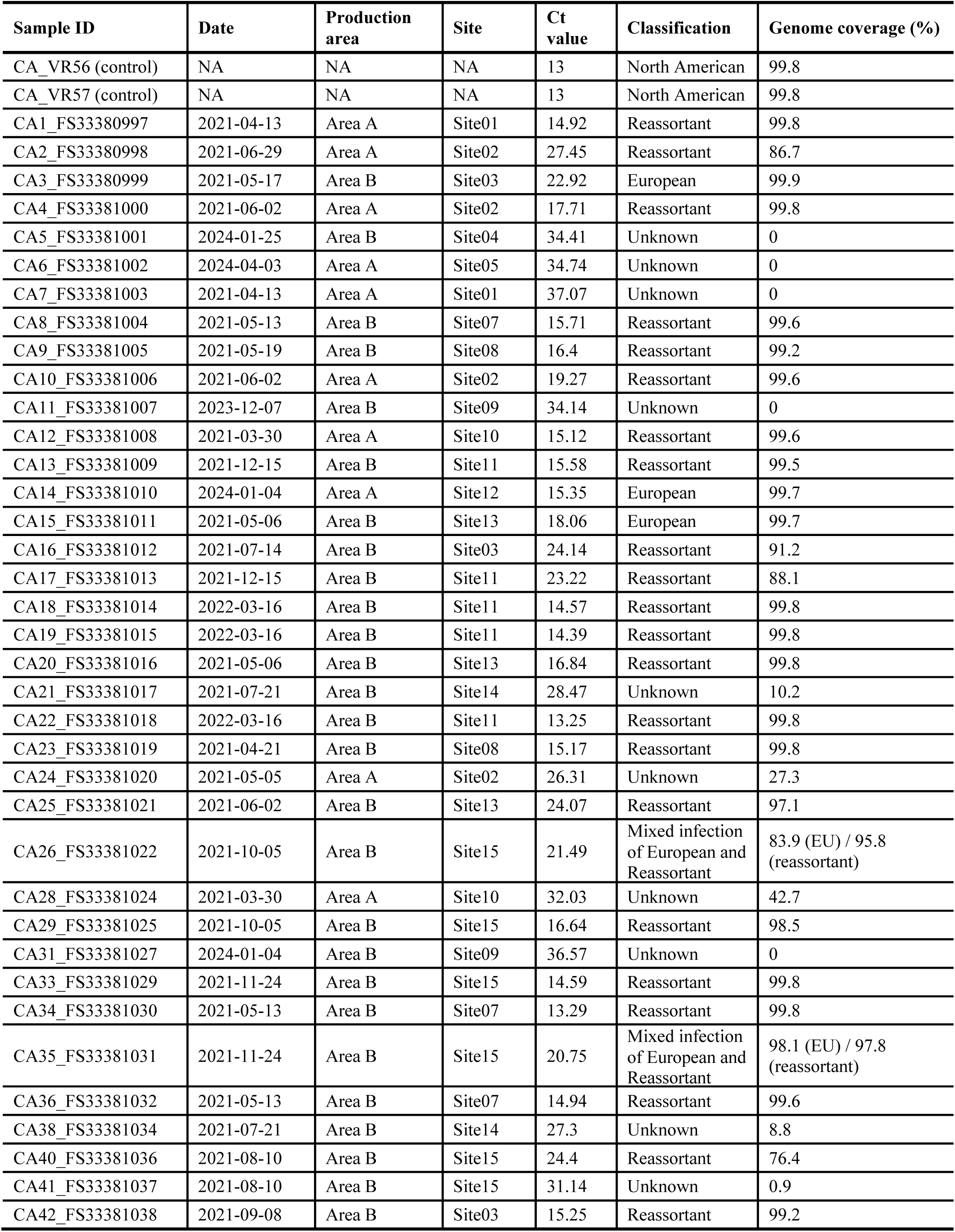
Metadata for the ISAV-positive samples included in this study. Sample information includes the sample identifier, collection date, anonymised production area and farm site, RT-qPCR Ct value, classification and genome coverage determined from whole-genome sequencing. Samples were classified as North American genotype, European genotype, reassortant, mixed infection or unknown. “Reassortant” denotes viruses containing European-genotype segment 3 and North American-genotype segments 1, 2 and 4–8, whereas “mixed infection” denotes samples containing both European-genotype and reassortant ISAV sequences. “Unknown” indicates that insufficient viral genomic sequence was recovered to determine the genotype, and “NA” indicates that metadata were not available.

| **Sample ID** | **Date** | **Production area** | **Site** | **Ct value** | **Classification** | **Genome coverage (%)** |
|---|---|---|---|---|---|---|
| CA_VR56 (control) | NA | NA | NA | 13 | North American | 99.8 |
| CA_VR57 (control) | NA | NA | NA | 13 | North American | 99.8 |
| CA1_FS33380997 | 2021-04-13 | Area A | Site01 | 14.92 | Reassortant | 99.8 |
| CA2_FS33380998 | 2021-06-29 | Area A | Site02 | 27.45 | Reassortant | 86.7 |
| CA3_FS33380999 | 2021-05-17 | Area B | Site03 | 22.92 | European | 99.9 |
| CA4_FS33381000 | 2021-06-02 | Area A | Site02 | 17.71 | Reassortant | 99.8 |
| CA5_FS33381001 | 2024-01-25 | Area B | Site04 | 34.41 | Unknown | 0 |
| CA6_FS33381002 | 2024-04-03 | Area A | Site05 | 34.74 | Unknown | 0 |
| CA7_FS33381003 | 2021-04-13 | Area A | Site01 | 37.07 | Unknown | 0 |
| CA8_FS33381004 | 2021-05-13 | Area B | Site07 | 15.71 | Reassortant | 99.6 |
| CA9_FS33381005 | 2021-05-19 | Area B | Site08 | 16.4 | Reassortant | 99.2 |
| CA10_FS33381006 | 2021-06-02 | Area A | Site02 | 19.27 | Reassortant | 99.6 |
| CA11_FS33381007 | 2023-12-07 | Area B | Site09 | 34.14 | Unknown | 0 |
| CA12_FS33381008 | 2021-03-30 | Area A | Site10 | 15.12 | Reassortant | 99.6 |
| CA13_FS33381009 | 2021-12-15 | Area B | Site11 | 15.58 | Reassortant | 99.5 |
| CA14_FS33381010 | 2024-01-04 | Area A | Site12 | 15.35 | European | 99.7 |
| CA15_FS33381011 | 2021-05-06 | Area B | Site13 | 18.06 | European | 99.7 |
| CA16_FS33381012 | 2021-07-14 | Area B | Site03 | 24.14 | Reassortant | 91.2 |
| CA17_FS33381013 | 2021-12-15 | Area B | Site11 | 23.22 | Reassortant | 88.1 |
| CA18_FS33381014 | 2022-03-16 | Area B | Site11 | 14.57 | Reassortant | 99.8 |
| CA19_FS33381015 | 2022-03-16 | Area B | Site11 | 14.39 | Reassortant | 99.8 |
| CA20_FS33381016 | 2021-05-06 | Area B | Site13 | 16.84 | Reassortant | 99.8 |
| CA21_FS33381017 | 2021-07-21 | Area B | Site14 | 28.47 | Unknown | 10.2 |
| CA22_FS33381018 | 2022-03-16 | Area B | Site11 | 13.25 | Reassortant | 99.8 |
| CA23_FS33381019 | 2021-04-21 | Area B | Site08 | 15.17 | Reassortant | 99.8 |
| CA24_FS33381020 | 2021-05-05 | Area A | Site02 | 26.31 | Unknown | 27.3 |
| CA25_FS33381021 | 2021-06-02 | Area B | Site13 | 24.07 | Reassortant | 97.1 |
| CA26_FS33381022 | 2021-10-05 | Area B | Site15 | 21.49 | Mixed infection of European and Reassortant | 83.9 (EU) / 95.8 (reassortant) |
| CA28_FS33381024 | 2021-03-30 | Area A | Site10 | 32.03 | Unknown | 42.7 |
| CA29_FS33381025 | 2021-10-05 | Area B | Site15 | 16.64 | Reassortant | 98.5 |
| CA31_FS33381027 | 2024-01-04 | Area B | Site09 | 36.57 | Unknown | 0 |
| CA33_FS33381029 | 2021-11-24 | Area B | Site15 | 14.59 | Reassortant | 99.8 |
| CA34_FS33381030 | 2021-05-13 | Area B | Site07 | 13.29 | Reassortant | 99.8 |
| CA35_FS33381031 | 2021-11-24 | Area B | Site15 | 20.75 | Mixed infection of European and Reassortant | 98.1 (EU) / 97.8 (reassortant) |
| CA36_FS33381032 | 2021-05-13 | Area B | Site07 | 14.94 | Reassortant | 99.6 |
| CA38_FS33381034 | 2021-07-21 | Area B | Site14 | 27.3 | Unknown | 8.8 |
| CA40_FS33381036 | 2021-08-10 | Area B | Site15 | 24.4 | Reassortant | 76.4 |
| CA41_FS33381037 | 2021-08-10 | Area B | Site15 | 31.14 | Unknown | 0.9 |
| CA42_FS33381038 | 2021-09-08 | Area B | Site03 | 15.25 | Reassortant | 99.2 |

### 2.3 Preparation of DNA libraries

DNA libraries were prepared from total RNA. First, a master mix containing 1 µL of dNTP mix (10 mM) and 1 µL of random hexamers was added to 10 µL of each RNA sample. Samples were incubated at 65 °C for 5 min and cooled on ice to bind hexamers. Following addition of 8 uL SuperScript III (SSIII) master mix containing 5× RT buffer, SSIII enzyme, RNaseOUT and DTT, the sample was incubated at 25 °C for 10 min, 55 °C for 60 min and 70 °C for 15 min to complete reverse transcription. Second-strand cDNA synthesis was performed using the NEBNext Ultra II Non-Directional RNA Second Strand Synthesis Module (New England Biolabs). Sixty microlitres of master mix containing 10× second-strand buffer, enzyme and nuclease-free water were added to each 20-µL first-strand cDNA reaction, followed by incubation at 16 °C for 2.5 h. Double-stranded cDNA was purified using 1× AMPure XP beads, eluted in 27 µL of Tris buffer (pH 8.0) and quantified using the Qubit dsDNA High Sensitivity Assay Kit. For end repair and A-tailing, 25 µL of cDNA was combined with 5 µL of end-repair/A-tailing master mix from the KAPA HyperPrep Kit (Roche) and incubated at 20 °C for 30 min and 65 °C for 30 min. Ligation was performed by adding 20 µL of ligation master mix and 5 µL of diluted KAPA universal adapter, followed by incubation at 20 °C for 60 min. Ligated DNA was purified using 0.8× AMPure XP beads and eluted in 24 µL of 10 mM Tris-HCl (pH 8.0). PCR amplification was performed in a total reaction volume of 50 µL, comprising 22.5 µL of the eluted DNA, 2.5 µL of ONT hybrid indices (10 µM), and 25 µL of KAPA HiFi HotStart ReadyMix (Roche). Cycling conditions comprised an initial denaturation at 98 °C for 45 s, followed by amplification cycles at 98 °C for 15 s, 65 °C for 30 s and 72 °C for 30 s, and a final extension at 72 °C for 1 min. The number of amplification cycles (1–19) was adjusted according to the input DNA concentration. PCR products were purified using 0.9× AMPure XP beads and eluted in 20 µL of nuclease-free water. DNA concentration was measured using a Qubit fluorometer, and fragment-size distributions were assessed using a TapeStation.

### 2.4 Probe design

To represent the genomic diversity of both ISAV genotypes, all 2,261 publicly available ISAV nucleotide sequences were downloaded from GenBank on 20 March 2023 and used to design a probe-capture hybridisation panel. Candidate 120-mer probes were designed to provide coverage across all eight ISAV genome segments, allowing up to five mismatches between each probe and its corresponding target sequence in accordance with the Twist Bioscience design criteria. Probes containing two or more ambiguous bases (N), or low-complexity sequence across at least 50% of their length, were excluded. The capture panel also contained probes targeting piscine myocarditis virus (PMCV); however, PMCV detection and sequence analysis were outside the scope of the present study, and only ISAV-derived sequencing data were analysed and reported. The final combined panel comprised 1,413 unique 120-mer probes.

### 2.5 Target enrichment

Samples were pooled together prior to enrichment. Each enrichment pool contained 10 indexed samples, with each contributing 150 ng of DNA, for a total pool mass of 1.5 µg. Sample volumes were calculated from individual DNA concentrations to ensure equal-mass pooling. The pooled library was purified using 1.8× KAPA Pure beads and eluted in 12 µL of blocking mixture containing 3 µL of 1 mM ONT hybrid blocker, 5 µL of Salmon Hybloc DNA (Applied Genetics Laboratories) and 4 µL of nuclease-free water.

Hybridisation of probes and post-capture amplification of samples to enrich for ISAV were performed using a modified Twist Target Enrichment protocol. A probe master mix containing 20 µL of hybridisation mix, 4 µL of the ISAV probe panel and 4 µL of nuclease-free water was denatured at 95 °C for 2 min and cooled on ice. The pooled library was heated at 95 °C for 5 min and placed on a magnetic rack. After the probe mixture had equilibrated to room temperature, the resuspended library was transferred to the probe mixture, vortexed and briefly centrifuged, and 30 µL of hybridisation enhancer was added. The reaction was incubated at 70 °C for 16 h.

Following hybridisation, the 70-µL capture reaction was transferred to 200 µL of washed streptavidin beads from the Twist Binding and Purification Beads Kit. The mixture was incubated on a shaker at room temperature for 30 min and washed according to the Twist Hybridisation and Wash Kit protocol. Post-capture PCR amplification of captured ISAV fragments was performed using 2.5 µL of ILMN primers and 25 µL of Equinox Library Amplification Mix. PCR was conducted in the presence of the streptavidin beads, with initial denaturation at 98 °C for 45 s; 5–15 cycles at 98 °C for 15 s, 60 °C for 30 s and 72 °C for 30 s; and a final extension at 72 °C for 1 min. Amplified products were purified using 0.9× AMPure XP beads, eluted in 15 µL of Tris buffer (pH 8.0), and assessed using Qubit and TapeStation assays.

### 2.6 Nanopore sequencing

Final enriched libraries were prepared for nanopore sequencing according to Oxford Nanopore Technologies (ONT) protocols. Approximately 400 fmol of enriched library DNA per reaction was adjusted to a total volume of 47 µL. Seven microlitres of NEBNext Ultra II End Prep Buffer, 3 µL of nuclease-free water and 3 µL of NEBNext Ultra II End Prep Enzyme were added sequentially, with gentle pipette mixing after each addition. The reaction was incubated at 20 °C for 20 min and 65 °C for 10 min and then held at 4 °C. DNA was purified using 0.4× AMPure XP beads and quantified using Qubit reagents. On the day of sequencing, adapter ligation was performed using the NEBNext Quick Ligation Kit. Briefly, 60 µL of DNA, 25 µL of Ligation Buffer, 10 µL of NEBNext Quick T4 DNA Ligase and 5 µL of sequencing adapter were combined in a LoBind tube, mixed gently and incubated at room temperature for 20 min. Ligated DNA was purified using 0.4× AMPure XP beads and quantified using Qubit. A total of 200 fmol of each library was loaded onto an R10.4.1 MinION flow cell and sequenced using a MinION Mk1B device. Sequencing-adapter ligation and loading followed the Ligation Sequencing Kit V14 (SQK-LSK114).

### 2.7 Sequence assembly and alignment

Basecalling was performed using Guppy v6.1.5 with the super-accuracy configuration for R10.4.1/E8.2 chemistry at 400 bases per second (dna_r10.4.1_e8.2_400bps_sup.cfg; Oxford Nanopore Technologies). Reads were demultiplexed based on ONT hybrid indices, and barcode and adapter sequences were trimmed using Porechop v0.2.4 with the --require_two_barcodes option, requiring the corresponding barcode at both ends of a read (Wick, 2018). For each genomic segment, reads were aligned simultaneously to a combined reference set comprising the corresponding European genotype (GenBank accession numbers HQ259671–HQ259678) and North American genotype (KU587556–KU587563) sequences using Minimap2 v2.24 (Li, 2018). Alignments were sorted and indexed using SAMtools v1.17 (Danecek et al., 2021), and an initial consensus was generated using samtools consensus. Consensus sequences were polished using Medaka v1.11.3 with the r1041_e82_400bps_sup_g615 model (Oxford Nanopore Technologies, 2023). Positions with <30× read depth were masked with N. Genome coverage was calculated as the proportion of nucleotide positions across the expected eight-segment genome represented at ≥30× depth, and consensus sequences with >70% coverage for each segment were retained for further analyses.

Potential mixed infections were inferred when sequenced genetic diversity within a single sample supported the presence of more than one variant. For segments containing different genotypes, separate consensus sequences were generated by mapping to the corresponding European and North American genotype references. To resolve distinct variants within the same genotype, candidate mixed single-nucleotide variants (SNVs) were identified directly from reference-mapped reads using a custom Python script implemented with pysam. Candidate sites were required to have ≥30× depth, ≥15% minor-allele frequency and at least five reads supporting the minor allele and were subsequently phased based on read-level allelic linkage across overlapping reads. Following read assignment, lineage-specific consensus sequences were generated with Medaka (Oxford Nanopore Technologies, 2023), with positions having <30× coverage in the corresponding variant alignment masked with N.

Nucleotide and corresponding amino-acid sequences generated in this study, together with sequences retrieved from GenBank on 29 October 2025, were aligned using MUSCLE v5.1 with the PPP algorithm and default parameters, as implemented in Geneious Prime 2024.0.5 (Edgar, 2022; Kearse et al., 2012). For phylogenetic analysis of segments 5 and 6, regions subject to frequent insertions or deletions were removed from the alignment. For segment 5, this included F protein insertions between amino acid residues 268 and 269 of the HPR0 reference sequence XCI12368. For segment 6, the HE HPR region spanning amino acid residues 337– 372 of the HPR0 reference sequence XCI12379 was removed. These regions were analysed separately to describe patterns of F protein insertion and HE protein HPR deletions.

### 2.8 Maximum-likelihood phylogenetic analysis

Maximum-likelihood (ML) trees were reconstructed separately for each ISAV genome segment using IQ-TREE 2 v2.3.6 (Minh et al., 2020). Each dataset comprised sequences generated in this study together with European and North American genotype sequences retrieved from GenBank on 29 October 2025. Only sequences for which both the country and year of sampling were available and that covered >70% of the corresponding genome segment were included. ModelFinder, implemented in IQ-TREE 2, was used to select the best-fitting nucleotide-substitution model for each dataset (Kalyaanamoorthy et al., 2017). Branch support was assessed using 1,000 ultrafast bootstrap replicates (Hoang et al., 2018). Trees were visualised and annotated with metadata using FigTree v1.4.4 (Rambaut, 2018) and the ggtree package in R (Yu et al., 2017). Trees were midpoint-rooted for visualisation, and established EU- and NA-genotype reference clades were labelled to support segment-specific genotype assignment, enabling detection of intergenotypic reassortant viruses.

### 2.9 Time-scaled phylogenetic analysis

To assess whether the dataset contained sufficient temporal signal for molecular-clock analysis, a maximum-likelihood tree was reconstructed using IQ-TREE 2 v2.3.6 without enforcing a molecular-clock model as described above. Root-to-tip genetic distances were then regressed against sampling dates using TempEst v1.5.3 (Rambaut et al., 2016).

Following assessment of the temporal signal, a time-scaled phylogeny of EU-genotype ISAV segment 3 and 5 sequences was reconstructed using BEAST v1.10.4 (Drummond and Rambaut, 2007; Suchard et al., 2018). ModelFinder selected TIM+F+G4 (segment 3) and TVM+F+R2 (segment 5) under the Bayesian information criterion. Because TIM/TVM is not implemented in BEAST v1.10.4, analyses used the more general GTR model with empirical nucleotide frequencies and gamma-distributed among-site rate heterogeneity approximated using four discrete categories (GTR+F+Γ4) (Tavaré, 1986; Yang, 1994). An uncorrelated lognormal relaxed clock and a strict molecular clock (Drummond et al., 2006) were each evaluated in combination with a constant population-size coalescent model (Griffiths and Tavaré, 1994; Kingman, 1982) and a Bayesian Skygrid model (Gill et al., 2013). For model comparison, each of the four model combinations was run in duplicate for 100 million MCMC steps, with samples recorded every 10,000 steps. Marginal- likelihood estimates obtained using path sampling and stepping-stone sampling identified the uncorrelated lognormal relaxed clock with a Bayesian Skygrid coalescent prior as the best-supported model (Lartillot and Philippe, 2006; Fan et al., 2011; Xie et al., 2011). Under this model, two independent MCMC chains were run for 250 million steps each and sampled every 25,000 steps. Convergence and mixing were assessed in Tracer v1.7.2 by inspecting parameter traces and effective sample sizes after combining the log files from the independent runs (Rambaut et al., 2018). A maximum clade credibility (MCC) tree was generated using TreeAnnotator v1.10.4 after discarding the first 10% of posterior trees as burn-in. The tree was visualised and annotated using ggtree with R version 4.5.2 (Yu et al., 2017).

Sequence alignments for all eight genome segments from this study were concatenated in the order of segments 1–8 and screened for evidence of intra-clade reassortment using RDP5 (Martin et al., 2021). As no evidence of intra-clade reassortment was detected, the concatenated alignment was subsequently used to reconstruct a time-scaled phylogeny in BEAST v1.10.4. The analysis used an HKY substitution model with empirical nucleotide frequencies, an uncorrelated lognormal relaxed molecular clock, and a constant population-size coalescent prior, which together represented the best-supported model combination.

### 2.10 Analysis of genome recovery in relation to Ct value

The association between diagnostic RT-qPCR Ct value and genome coverage was assessed using Spearman’s rank correlation (Spearman, 1904). For visualisation of the non-linear relationship between Ct value and genome coverage, a four-parameter logistic decay model was fitted to the observed data using nonlinear least squares (Bates and Watts, 1988). The model included upper and lower asymptotes, a midpoint and a slope parameter describing the decline in genome coverage with increasing Ct value. The fitted curve was used descriptively to illustrate the overall relationship between Ct value and genome recovery in this dataset and was not used to define a diagnostic or sequencing-success threshold. Data analysis was performed in R version 4.5.2, and visualisation was performed using ggplot2 with R version 4.5.2 (Wickham, 2016).

## 3. Results

### 3.1 Sequencing performance and genome recovery

A total of 37 ISAV-positive Canadian samples collected between 2021 and 2024 were selected for whole-genome sequencing. Diagnostic RT-qPCR Ct values ranged from 13.25 to 37.07. Genome coverage was significantly negatively correlated with Ct value (Spearman’s ρ = −0.854, P < 0.001), with genome recovery decreasing as Ct increased. Among the 19 samples with Ct values <20, all yielded >99% genome coverage. Two different genomes indicative of mixed infection were recovered from sample CA35, which had a Ct value of 20.75, with genome coverages of 98.1% and 97.8%, respectively. Similarly, two genomes were recovered from sample CA26, which had a Ct value of 21.49, with genome coverages of 83.9% and 95.8%, respectively. Among the five samples with Ct values between 22 and 25, genome coverage ranged from 99.8% to 76.4%. Genome recovery was efficient across 8 segments as well as for both European (EU) and North American (NA) genotypes (Figure S1). Samples with Ct values >26 generally yielded <50% genome coverage, and some yielded no recoverable sequence. One exception was a sample with a Ct value of 27.45 that yielded 86.7% genome coverage (Figure 3; Table 1). Most samples with Ct values >30 yielded no recoverable genome sequence. Of the five samples collected in 2023–2024, four had Ct values >34 and yielded insufficient sequence for genotype assignment; the remaining sample, collected in 2024, yielded 99.7% genome coverage.

**Figure 3.**
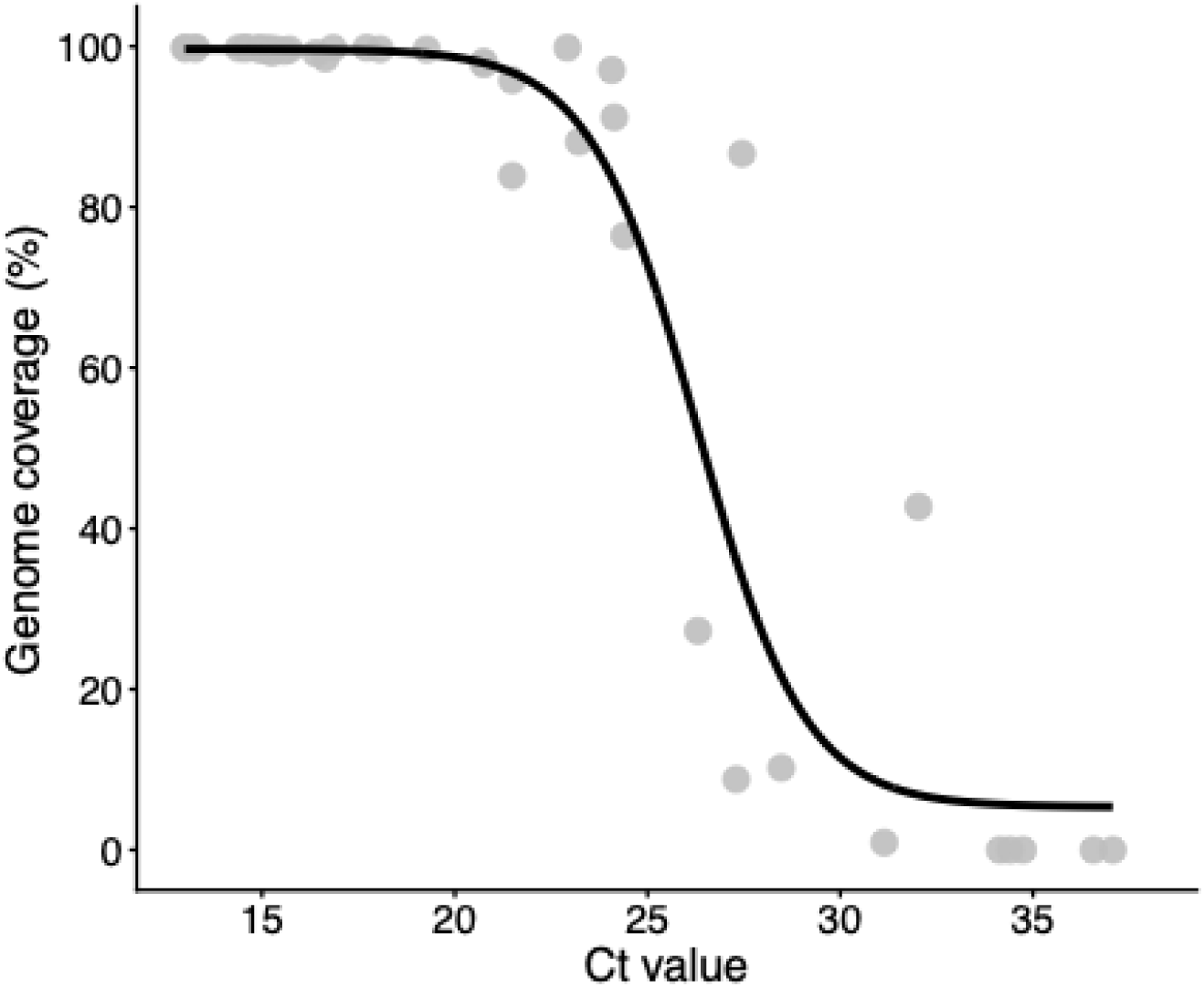
Relationship between diagnostic RT-qPCR Ct value and ISAV genome coverage. Genome coverage was calculated as the proportion of nucleotide positions across the expected eight-segment genome represented at ≥30× read depth. Samples with Ct values <25 generally yielded near-complete genomes, whereas recovery declined markedly at Ct values >25. The black line represents a four-parameter logistic decay curve fitted by nonlinear least squares, illustrating the overall decline in genome coverage with increasing Ct value.

Only consensus sequences for individual segments with >70% coverage were retained for downstream analyses. Twenty-seven Canadian samples yielded 29 genome assemblies because two samples contained mixed infections. These samples were derived from 10 farm sites across two Bay of Fundy production areas, with one to six genome assemblies obtained per farm. Two positive-control samples, VR56 and VR57, both with Ct values of approximately 13, yielded near-complete genomes. The final analysis dataset therefore comprised 31 consensus genomes: 29 from the Canadian samples and two from the positive controls.

### 3.2 ISAV genotypes identified by target-enrichment whole-genome sequencing

Whole-genome sequencing identified three infection patterns among samples: infection with an EU-genotype virus, infection with a reassortant virus containing segments of both NA and EU origin, and a mixed infection involving an EU-genotype virus and the identified reassortant virus (Figure S2). The reassortant virus contained EU-genotype segment 3 in an otherwise NA-genotype genomic background (Table 1, Figure 4, Figure S2). Twenty-four genomes were assigned to the reassortant virus, from nine farm sites: Sites 01, 02, 03, 07, 08, 10, 13 and 15 in 2021, and Site 11 in 2022.

**Figure 4.**
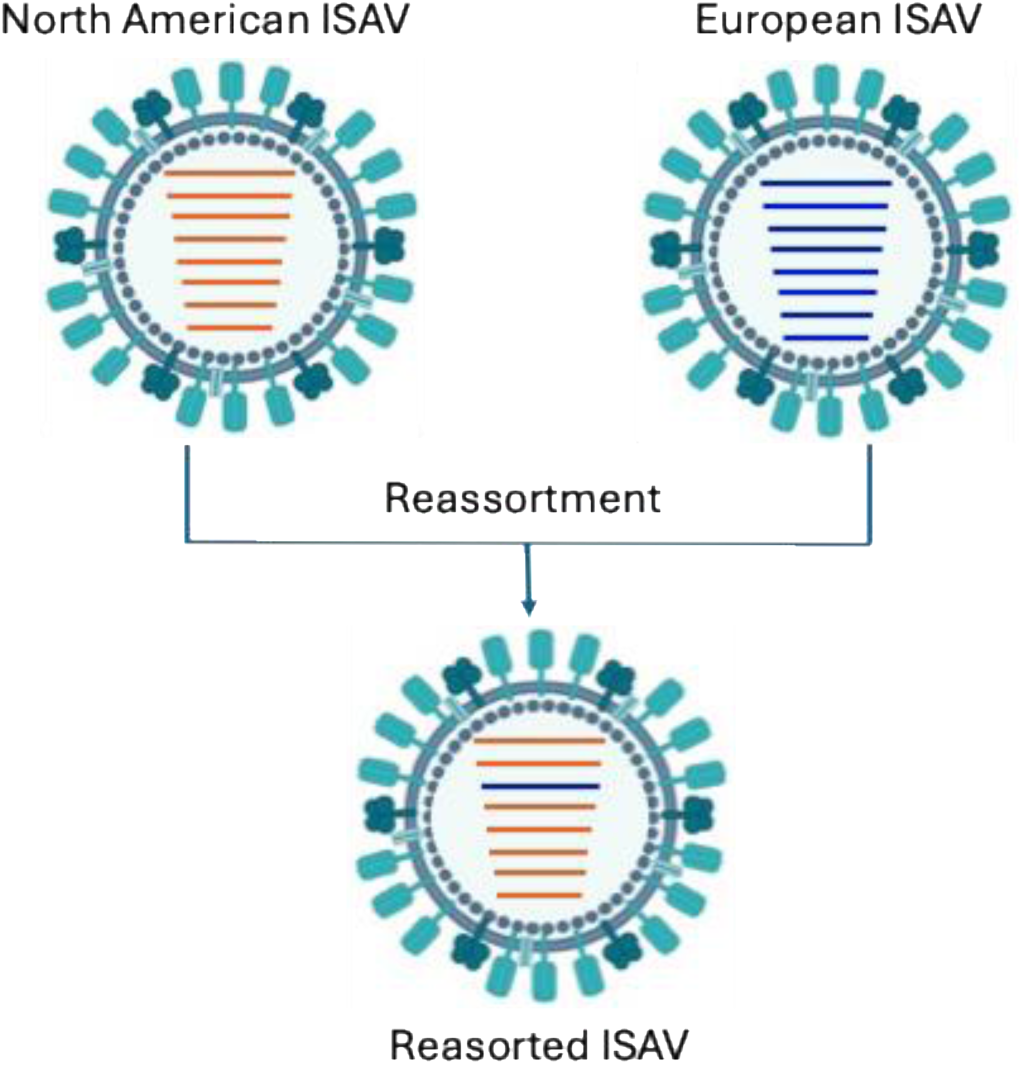
Schematic representation of reassortment between North American- and European-genotype ISAV. Red genome segments represent the North American genotype, and blue genome segments represent the European genotype. The reassortant ISAV contains European-genotype segment 3 together with North American-genotype segments 1–2 and 4–8.

Three samples, CA3, CA14 and CA15, contained non-reassortant EU-genotype viruses. They were collected from Sites 03 and 13 in 2021 and Site 12 in 2024. Two samples, CA26 and CA35, collected from Site 15 in 2021, contained mixed infections involving both an EU-genotype virus and the reassortant lineage, such that two distinct EU-genotype segment 3 sequences were recovered from each sample, alongside distinct NA and EU forms of all other segments. The co-infection was independently detected in two separate sequencing runs, with both samples sequenced twice and negative controls included in each run for confirmation. We attributed the segment 3 consensus sequence that showed high identity to segment 3 sequences from other Canadian reassortant viruses to the reassortant lineage, given similarity of the other NA segments identified. The other consensus sequence was attributed to the co-occurring EU-genotype virus segments, consistent with the genotype assignments of the other segments that did not show high identity to those from samples containing only the reassortant form. The two positive controls, VR56 and VR57, were classified as NA-genotype ISAVs (Table 1), confirming that the absence of an NA-genotype segment 3 was not attributable to a failure of the approach to detect this genotype.

Samples with insufficient sequence data for reliable genotype assignment were classified as unknown and excluded from further analyses (Table 1).

### 3.3 Phylogenetic evidence for the emergence and spread of a reassortant ISAV lineage

We first explored the evolutionary history of the EU-NA reassortant viruses to characterise the different evolutionary histories identifiable from each of the different segments. All reassortant virus segments clustered together as a clade in all ML trees with strong support (bootstrap ≥79%; Figure 5, Figures S3–S10). This shared genome constellation and consistent phylogenetic clustering across segment-specific trees are most consistent with descent from a single ancestral reassortant rather than repeated independent reassortment events. In segment 3, these reassortant viruses were most closely related to an EU-genotype HPR0 ISAV sampled in Hancock County, Maine (USA) in 2016 (bootstrap = 100%). The available genome segments from this virus consistently showed EU-genotype ancestry, with no evidence of reassortment between the European and North American genotypes. In all other segments, the reassortant viruses were most closely related to NA-genotype ISAVs previously sampled in Canada (bootstrap ≥79%), including both HPR0 and HPRdel collected in 2012 and 2013, which had no evidence of intergenotypic reassortment (Figure 5, Figures S3–S10).

**Figure 5.**
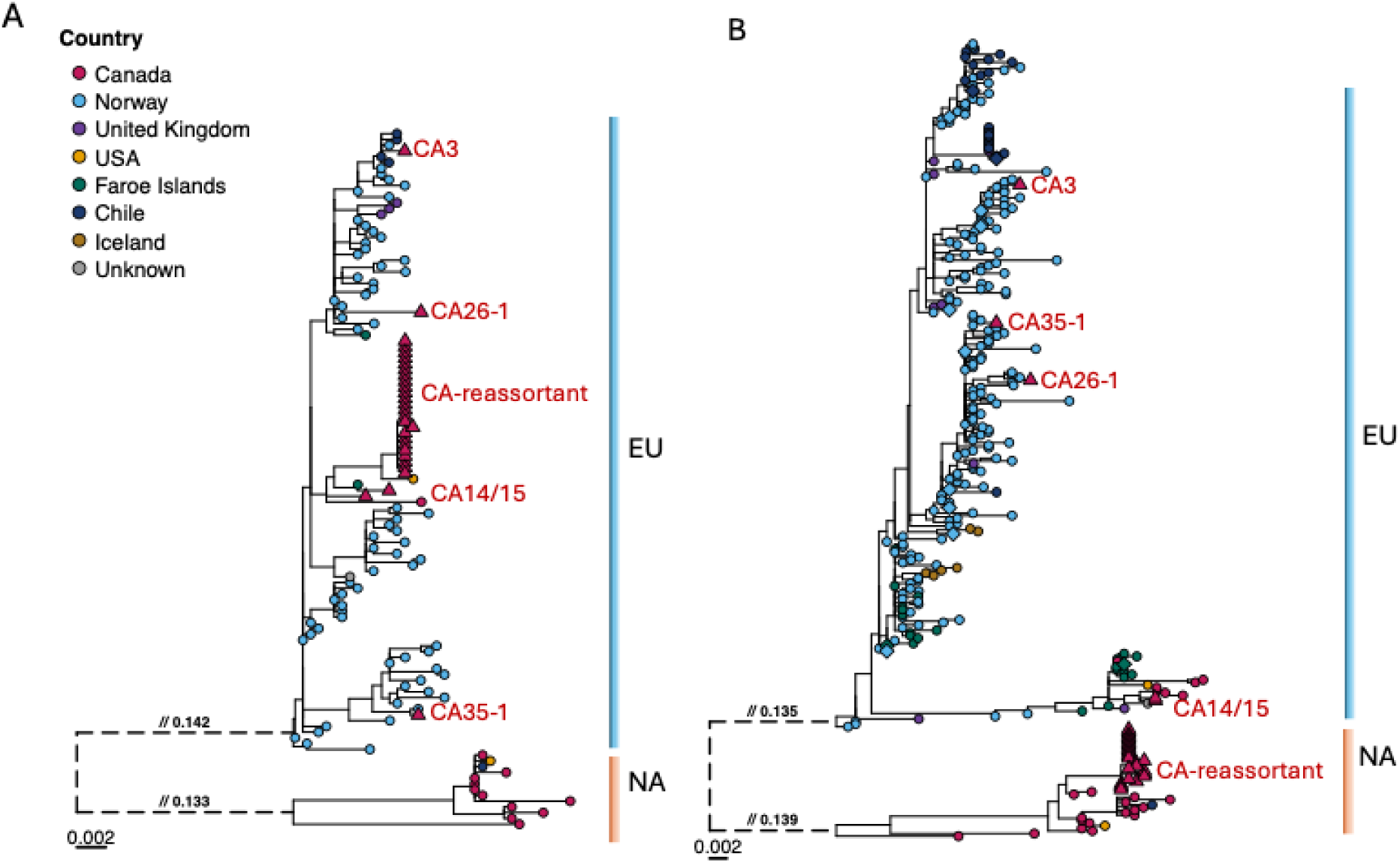
Maximum-likelihood phylogenies of ISAV segments 3 and 6. Trees were inferred from nucleotide alignments of segment 3 (A) and segment 6 (B). Tips are coloured by country of sampling. Canadian sequences generated in this study are shown as triangles and publicly available sequences as circles; Canadian sample labels are shown in red. In the segment 6 tree, country-specific clades containing >10 sequences were collapsed and are indicated by diamonds. Vertical bars denote the European (EU) and North American (NA) genotype clades. The reassortant lineage clusters within the EU genotype for segment 3 and within the NA genotype for segment 6. Exceptionally long branches were truncated for visualisation and are shown as dashed lines, with omitted distances labelled above the branches. Scale bars indicate 0.002 nucleotide substitutions per site. Bootstrap scores are not shown to aid visual clarity, but are presented in tree files in Files 3 and 6.

The ML trees of EU-genotype segment 3 sequences and concatenated genome sequences of reassortant viruses were separately rooted in TempEst to maximise the correlation between root-to-tip genetic distance and sampling date. The resulting regressions showed positive temporal associations for both datasets, with a correlation coefficient of 0.54 (R² = 0.29) for the complete EU-genotype segment 3 dataset and 0.50 (R² = 0.25) for the concatenated reassortant dataset, indicating moderate, although not strong, temporal signal (Figure S11A-B). The posterior mean tMRCA of the sampled Canadian reassortant cluster in the Segment 3 MCC tree was September 2016 (95% HPD: March 2013–June 2019). Analysis of the concatenated genomes of this clade yielded a broadly consistent estimate of July 2014 (95% HPD: July 2006–July 2020). Despite differences in the datasets and demographic models used, the overlapping 95% HPD intervals indicate that the two tMRCA estimates are concordant, and support the validity of using a molecular clock model to analyse this dataset. The common ancestor shared by this clade and its closest sampled segment 3 relative from Maine was dated to July 2011 (95% HPD: May 2006–September 2015). The posterior mean tMRCA of all EU-genotype segment 3 sequences included in the analysis was November 1956 (95% highest posterior density [HPD]: June 1933– December 1974). Segment 3 of the Canadian reassortant viruses formed a strongly supported monophyletic clade within a broader lineage containing ISAV sequences from Maine, the Faroe Islands and Canada (posterior support = 1.00; Figure 6A), consistent with a single reassortment event. The tMRCA of this clade indicates that this reassortant event likely occurred between July 2011 (95% HPD: May 2006–September 2015) and September 2016 (95% HPD: March 2013–June 2019).

**Figure 6.**
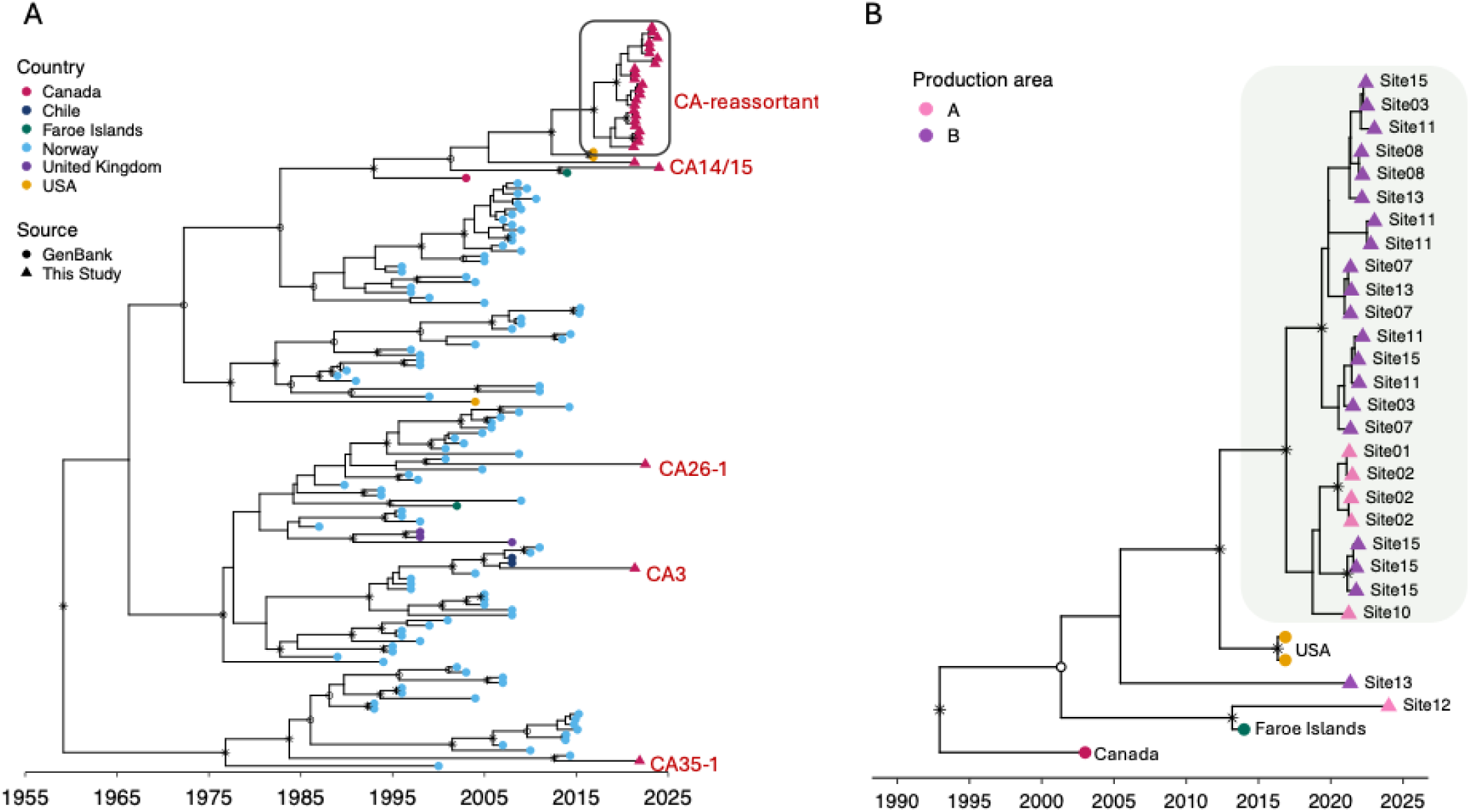
Time-scaled maximum clade credibility (MCC) phylogeny of European-genotype ISAV segment 3 sequences. (A) Segment 3 phylogeny with tips coloured by country of sampling. Publicly available GenBank sequences are shown as circles and Canadian sequences generated in this study as triangles; Canadian sample labels are shown in red. The outlined region identifies the clade containing the Canadian reassortant lineage. (B) Enlarged view of that clade. Canadian sequences are coloured by production area and labelled by farm site; publicly available sequences from Canada, the USA and the Faroe Islands are shown as coloured circles. The shaded region identifies the Canadian reassortant cluster. Asterisks indicate nodes with posterior probabilities >0.80, and circles indicate nodes with posterior probabilities >0.50 and ≤0.80. The horizontal axis is calendar time.

Reassortant viruses from nine farm sites across both production areas were interspersed throughout the clade in segment 3 (Figure 6B), indicating widespread transmission of the reassortant virus in the Bay of Fundy. Although some sequences from the same production area or farm site grouped together, including several viruses from Site 15 in Area B (posterior probability = 1) and viruses from Site 02 in Area A (posterior probability =1.00), posterior support for most internal nodes within the segment 3 clade was generally low, and the overall topology did not support segregation by farm site or sampling year (Figure 6B, Figure S12).

To determine whether additional phylogenetic resolution could be obtained from genome-wide sequence data, a time-scaled MCC tree was reconstructed from concatenated segment 1-8 sequences of reassortant viruses for which all eight genome segments were available. The concatenated analysis showed a similar pattern to that observed in phylogenies of segment 3, consistent with the lack of reassortment within this clade. Substantial clustering by farm site was evident, with several same-site groups strongly supported, including viruses from Sites 02, 13 and a subset of those from Site 15 (posterior probability = 1). The tree also showed pronounced structure by production area. Most viruses from Production Area B formed a strongly supported major clade (posterior probability = 1). The second major lineage (posterior probability = 0.93) contained the remaining Production Area B viruses together with several viruses from Production Area A. Thus, concatenation increased resolution of several internal relationships and revealed clearer geographic structuring by production area and farm site, although limited posterior support for some internal nodes restricted more detailed reconstruction of transmission routes among individual farms (Figure S13).

### 3.4 Phylogenetic diversity of non-reassortant EU-genotype ISAVs

Unlike the reassortant viruses, the five EU-genotype genomes identified in the Canadian samples—CA3, CA14, CA15, CA26-1 and CA35-1—did not form a single clade but occupied several distinct positions within the EU-genotype phylogenies (Figures S3–S10). CA3, CA26-1 and CA35-1 occupied separate clades dominated by Norwegian sequences in the segment 3-6 ML trees (bootstrap > 90%; Figure 5; Figures S5-8). CA26-1 and CA35-1 clustered closely within Norwegian ISAVs in the segment 1 and 2 ML phylogenies (bootstrap = 100% and 98%, respectively) and with Norwegian and Chilean ISAVs in segment 7 ML phylogeny, although with lower support (bootstrap = 64%; Figures S3, S4, S9).

The ML trees of the EU-genotype segment 5 sequences showed positive temporal signal, with correlation coefficients of 0.61 (R² = 38) (Figure S11C). Consistent with the ML analyses, CA3, CA26-1 and CA35-1 remained phylogenetically distinct in the segment 3 and 5 time-scaled phylogenies, and were separated from both the reassortant lineage and each other (Figure 6A; Figure 7). In the time-scaled phylogeny of segment 5, CA35-1 shared a recent common ancestor with a Norwegian ISAV sampled in 2024, with a tMRCA estimated at September 2020 (95% HPD: May 2018–October 2021; posterior probability = 1.00). CA3 also shared a recent common ancestor with a Norwegian ISAV sampled in 2024, with its tMRCA estimated at July 2020 (95% HPD: August 2018–April 2021; posterior probability = 0.94). CA26-1 shared a recent common ancestor with Norwegian ISAVs sampled in 2025, with the tMRCA estimated at approximately April 2019 (95% HPD: March 2016–May 2021; posterior probability = 0.94). CA3 and CA26-1 shared a more recent common ancestor dating to March 2010 (95% HPD: June 2005–August 2014; posterior probability = 0.88). The broader clade containing CA35-1, CA3 and CA26-1 had an estimated tMRCA of approximately July 1991 (95% HPD: February 1988– May 1994; posterior probability = 0.86) (Figure 7).

**Figure 7.**
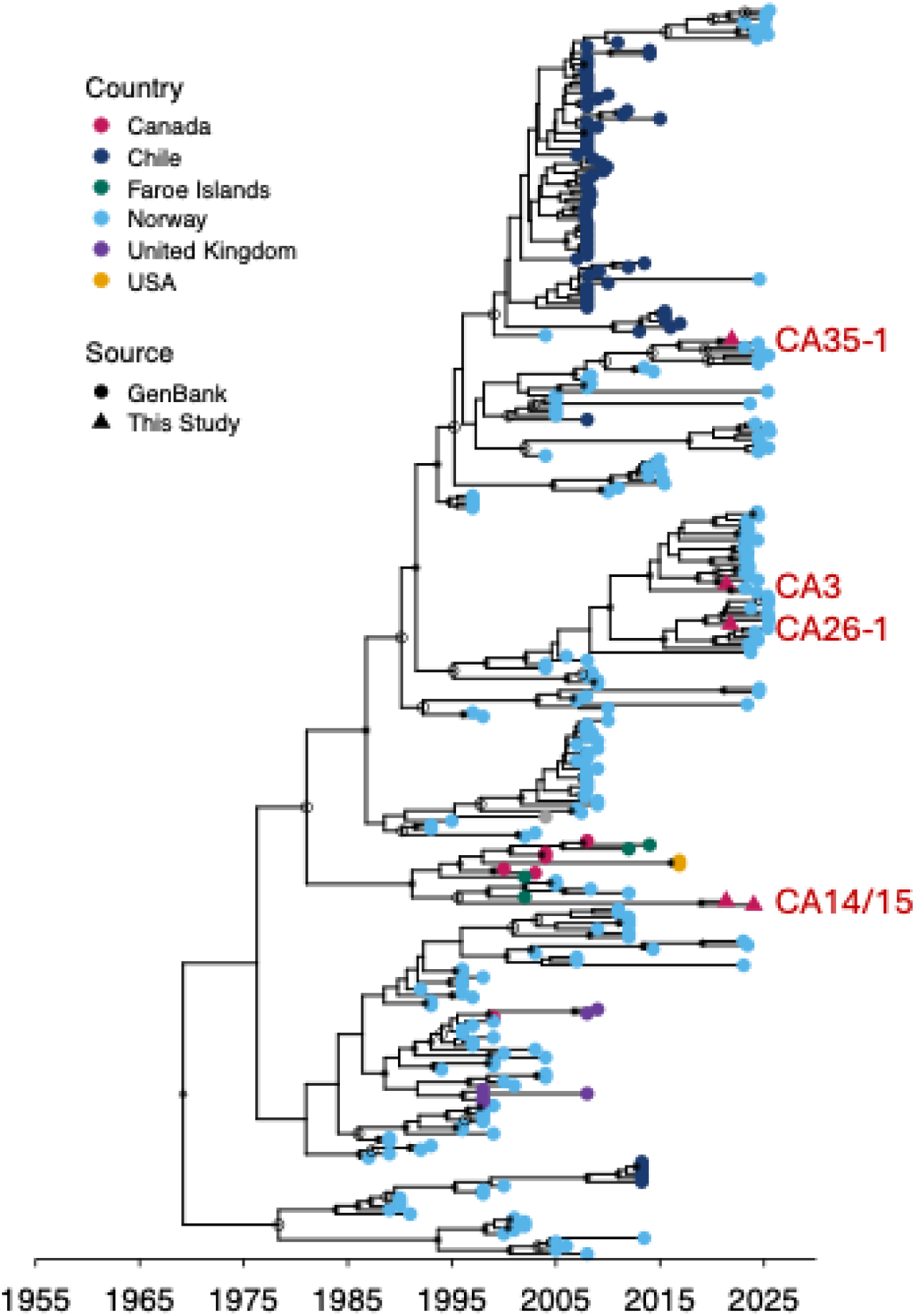
Time-scaled maximum clade credibility (MCC) phylogeny of European-genotype ISAV segment 5 sequences with tips coloured by country of sampling. Publicly available GenBank sequences are shown as circles and Canadian sequences generated in this study as triangles; Canadian sample labels are shown in red and labelled. Asterisks indicate nodes with posterior probabilities > 0.80, and circles indicate nodes with posterior probabilities > 0.50 and ≤ 0.80. The horizontal axis is calendar time.

CA14 and CA15, which originated from two different farm sites, formed a strongly supported sister clade in the segment 1, 4, 5 and 6 ML trees (bootstrap ≥ 98%). In segment 2 and 7 phylogenies, they did not form an exclusive monophylogenetic clade but occurred within the same broader clade alongside additional Canadian and non-Canadian ISAV sequences, including viruses from the Faroe Islands, the USA and Norway (bootstrap ≥ 96%; Figures S3–S9). In the segment 3 phylogenies, CA14 and CA15 were positioned near the base of the clade containing the Canadian reassortant viruses and clustered with previously reported EU-genotype ISAVs from Canada, the USA and the Faroe Islands (bootstrap = 96% in Figures 5A and posterior probability = 1 in 6A). Their segment 6 sequences also clustered within the EU genotype and were closely related to ISAVs previously detected in the Faroe Islands, the USA, the UK and Canada (bootstrap = 96%). They were clearly separated from the NA-genotype segment 6 sequences of the reassortant lineage (Figure 5B).

In the time-scaled phylogeny of segment 5, they share tMRCA at 2018.9593 (95% HPD: 2014.3733 - 2021.2081; posterior = 1). tMRCA of them with the most closely-related sequences from Faroe Islands and Norway was 1995.5839 (95% HPD: 1990.5773 – 1999.64; posterior = 1) (Figure 7; Figure S14).

### 3.5 Amino-acid variation in the HE and F proteins of Canadian ISAVs

Given their impact on pathogenicity, we characterised variation in the F protein amino-acid position 266 (segment 5) and HPRdel deletion sites (segment 6). Among the 28 Canadian genomes for which segment 5 met the coverage threshold (70%), 26 encoded leucine (L) at F-protein amino-acid position 266 and contained no insertion near the cleavage site. CA15 encoded arginine (R) at this position without an insertion, whereas CA35-1 encoded glutamine (Q) and contained an insertion (Figure 8).

**Figure 8.**
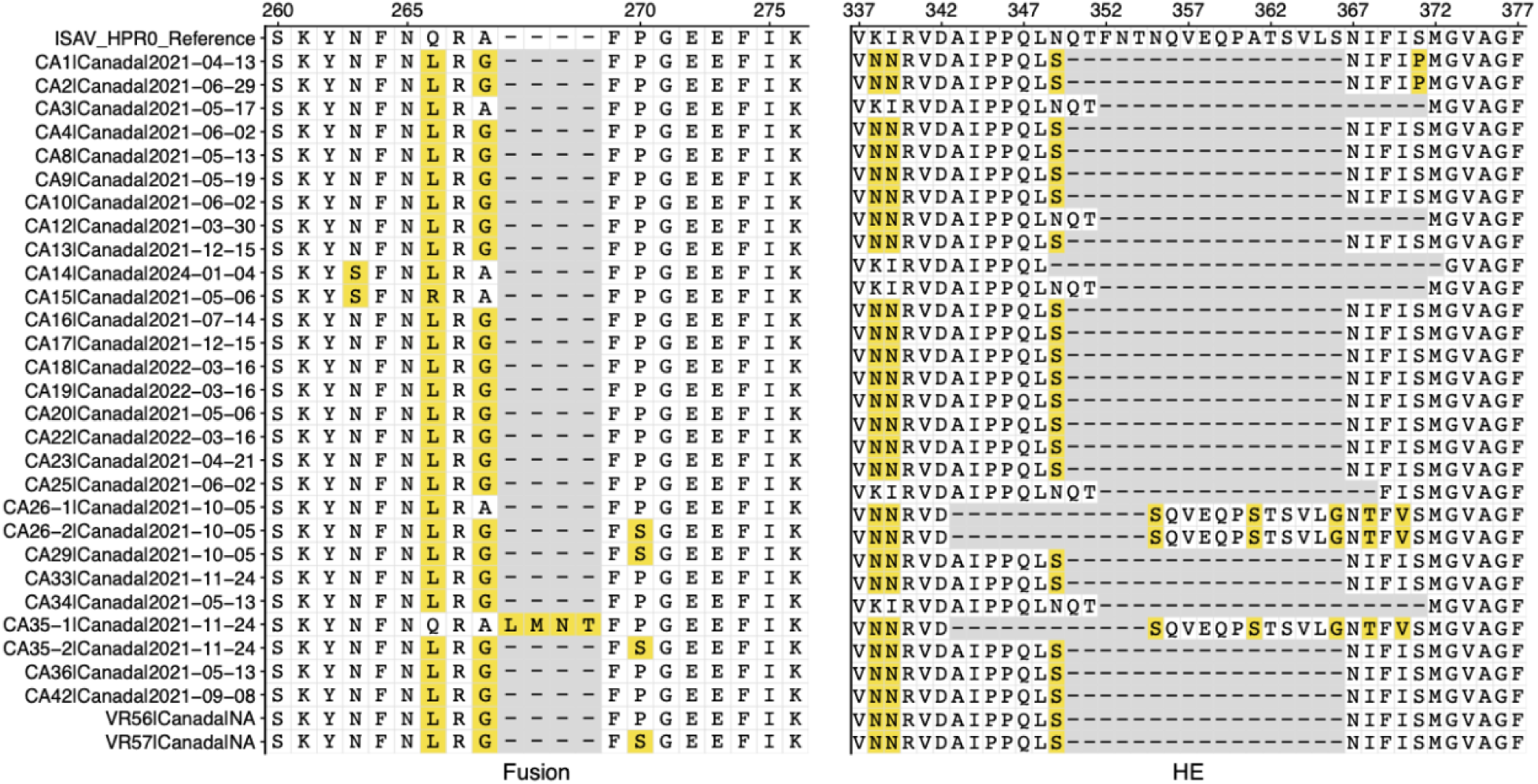
Amino-acid variation in the infectious salmon anaemia virus fusion (F) and haemagglutinin–esterase (HE) proteins. Amino-acid alignments are shown for the F-protein region containing residue 266 and the adjacent insertion site (left) and the HE highly polymorphic region (HPR; right). Norwegian HPR0 sequences XCI12368 and XCI12379 were used as the reference sequences for the F and HE alignments, respectively, and are labelled as ISAV_HPR0_Ref. Amino-acid positions are numbered relative to the corresponding HPR0 reference sequence. Yellow shading indicates residues that differ from the reference, and grey cells with dashes indicate alignment gaps. Sequence names are shown for the F-protein alignment and correspond to the same sequence rows in the HE alignment.

All viruses for which the segment 6 HPR was recovered were HPRdel variants (Figure 8). Relative to the HPR0 reference sequence (GenBank accession XCI12379), NA-genotype segment 6 sequences from the reassortant lineage displayed three deletion patterns: a 12-amino-acid deletion spanning aa 343–354, a 17-amino-acid deletion spanning aa 350–366 and a 20-amino-acid deletion spanning aa 352–371. EU-genotype segment 6 sequences from CA3, CA14, CA15, CA26-1 and CA35-1 also displayed three deletion patterns. Two were identical to patterns also observed in the reassortant viruses: a 17-amino-acid deletion spanning aa 352–368 and a 20-amino-acid deletion spanning aa 352–371. The third was a distinct 24-amino-acid deletion spanning aa 349–372 that was not identified amongst the studied EU-NA reassortant viruses.

## 4. Discussion

This study used target-enrichment whole-genome sequencing to characterise ISAV genomic diversity in farmed Atlantic salmon from the Bay of Fundy, New Brunswick. We identified 24 viruses carrying a European (EU)-genotype segment 3 within an otherwise North American (NA)-genotype genome. These viruses were detected at nine farm sites across two production areas during 2021 and 2022, indicating widespread circulation of the reassortment genotype in this region. All sequenced viruses were virulent HPRdel ISAV forms, but we acknowledge that our ability to sequence HPR0 was limited by the sensitivity of our sequencing, making it impossible to investigate HPR0 diversity in the Bay of Fundy. We also identified several phylogenetically distinct EU-genotype viruses and two mixed infections involving both an EU-genotype virus and the reassortant lineage. These findings reveal substantial ISAV genomic diversity across the sampled production areas and demonstrate that phylogenetic inference based on a single genome segment does not accurately represent the genomic ancestry or genomic diversity of circulating viruses.

### 4.1 Intergenotypic ISAV reassortant lineage

Reassortment has previously contributed to the emergence and diversification of ISAV (Devold et al., 2006; Plarre et al., 2012), most notably during the Chilean ISA epidemic in which circulating viruses contained segments derived from distinct EU-genotype ancestors (Cottet et al., 2010). To our knowledge, the Canadian reassortant lineage identified here provides the first documented evidence of reassortment between the European and North American ISAV genotypes. The shared genomic constellation and monophyletic clustering of the 24 reassortant viruses in all segmental phylogenies are most consistent with a single ancestral reassortment event followed by onward transmission of the reassortant lineage. Although repeated reassortment events involving closely related parental viruses could theoretically generate the same genomic constellation, this scenario would require multiple independent coinfections involving genetically similar parental viruses to repeatedly produce the same segment combination and is therefore less likely.

The segment 3 time-scaled phylogeny dated the posterior mean tMRCA of the sampled reassortant lineage to approximately September 2016, around five years before the first sampling event in 2021. Under a single-reassortment scenario, the reassortment event would have occurred along the ancestral branch between the divergence of the EU-derived segment 3 lineage from its closest sampled relative in Hancock County, Maine, USA (only 200 km away), and the diversification of the sampled reassortant clade. The outer bounds of the corresponding 95% HPD intervals define an approximate reassortment window of May 2006–June 2019. However, this range is not a formal posterior interval for the reassortment event and should be interpreted cautiously in light of the moderate temporal signal and limited historical sampling of ISAV. Nevertheless, our analyses suggest that the reassortant virus has almost certainly been circulating for multiple years.

In other members of the family *Orthomyxoviridae*, reassortment can reduce fitness by disrupting co-adapted molecular interactions required for efficient replication and transmission, but it can also generate variants that outcompete previously circulating viruses (McDonald et al., 2016). The reassortant viruses retained the ISAV virulence-associated genes in segment 5, encoding the F protein, and segment 6, encoding the HE protein, from the NA genotype (Markussen et al., 2008; Cárdenas et al., 2022). In contrast, segment 3, encoding the nucleoprotein (NP), was derived from the EU genotype. NP is an essential component of viral ribonucleoprotein complexes involved in transcription, genome replication and packaging (Goić et al., 2008; Zheng et al., 2013). In influenza A viruses, reassortment involving NP can influence host adaptation, transmissibility and pathogenicity (Wasilenko et al., 2008; Ince et al., 2013; Hao et al., 2016; Thomas et al., 2024), suggesting that replacement of this segment could have biological consequences beyond recognised ISAV virulence markers. Whether acquisition of the EU-genotype NP alters ISAV phenotype will require experimental investigation. Nevertheless, detection of the reassortant lineage at multiple farms and across two sampling years indicates that this genome constellation was compatible with onward transmission and persistence rather than representing a transient reassortment event. The lineage was not recovered from samples collected in 2023 or 2024; however, four of the five samples from those years had Ct values >34 and yielded insufficient sequence for genotype assignment. Its subsequent circulation therefore remains unresolved, and broader contemporary sampling will be required to determine whether the lineage persists in the region or elsewhere in Canada, the USA or globally.

The occurrence of closely related viruses at different sites is consistent with regional circulation and epidemiological connectivity among farms, although phylogenetic data alone cannot identify the underlying transmission pathways. Possible routes include transfers of infected fish, movements of shared vessels or equipment, waterborne transmission and transmission through unsampled hosts or reservoirs, including wild salmon (Plarre et al., 2005; Aldrin et al., 2010; Mardones et al., 2014; Gagné and LeBlanc, 2018; Gautam et al., 2018; Duault et al., 2025; Zhao et al., 2026). Integration of genomic data with stocking records, fish and vessel movements, outbreak timing and hydrodynamic connectivity would be required to distinguish among these routes.

### 4.2 Multiple EU-genotype ISAV lineages in Canada

Whilst no non-reassortant NA-genotype ISAVs were identified, we sequenced five EU-genotype ISAVs: three in the apparent absence of the reassortant virus (CA3, CA14, CA15), and two (CA26-1 and CA35-1) as co-infections alongside the reassortant. The five non-reassortant EU-genotype viruses did not form a single Canadian clade. Instead, their positions across most segment phylogenies were consistent with at least four distinct phylogenetic lineages, three represented individually by CA3, CA26-1 and CA35-1 and a fourth represented by CA14 and CA15. Overall, the phylogenetic pattern in which viruses from Canada are interspersed with those from other locations likely implies multiple introductions of the EU genotype ISAV into Canada, as previously suggested for Atlantic Canada (Gagné and LeBlanc, 2018).

CA14 and CA15 were more closely related to EU-genotype ISAV previously detected in the USA, Canada and the Faroe Islands. They clustered closely together in most segment phylogenies despite being sampled nearly three years apart, suggesting persistence of a related EU-genotype lineage in the region. However, their discordant placements in segments 2 and 7 were strongly supported (bootstrap > 90%), likely indicating segment-specific phylogenetic incongruence consistent with reassortment among EU-genotype lineages.

CA3, CA26-1 and CA35-1 consistently clustered with Norwegian-dominated EU-genotype lineages across multiple genome segments, with these relationships strongly supported in several phylogenies, although their phylogenetic placement varied amongst segments. The time-scaled segment 5 phylogenies further showed relatively recent shared ancestry, around 2020, between each of these Canadian viruses and Norwegian ISAVs, sampled in 2024 or 2025. In contrast, the common ancestor of CA3, CA26-1 and CA35-1 was estimated to date to approximately the 1990s. These patterns indicate that the three viruses represent distinct and divergent EU-genotype lineages rather than descendants of a single recent Canadian lineage. Their recent common ancestry with Norwegian viruses is compatible with relatively recent trans-Atlantic connectivity of these lineages, although it does not establish Norway as the direct geographic source. ISAV exchange between European and North American wild Atlantic salmon at North Atlantic feeding grounds has previously been proposed as one possible mechanism linking viruses across the North Atlantic (Gagné and LeBlanc, 2018). The observed phylogenetic pattern could therefore reflect multiple introductions into Canada, persistence and diversification of previously introduced EU-genotype lineages, transmission through unsampled intermediate populations, or a combination of these processes. More geographically balanced and temporally continuous whole-genome sampling of ISAV from farmed and wild Atlantic salmon across the North Atlantic, combined with information on movements of live fish and other relevant aquaculture pathways, will be required to resolve the origins and routes of trans-Atlantic ISAV dispersal.

Previous analyses also indicate that the Canadian ISAV population comprises endemic North American-genotype viruses together with several incursions of EU-genotype viruses (Gagné and LeBlanc, 2018). Interestingly, although NA-genotype ISAV has previously been detected in the Bay of Fundy, including isolates sampled in the late 1990s (Kibenge et al., 2007b; Kulshreshtha et al., 2010), we did not identify any evidence of a wholly NA-genotype virus circulating in the Bay of Fundy, New Brunswick region. This absence should not be interpreted as evidence that NA-genotype viruses are no longer circulating in the region, given the spatial and temporal limitations of our sampling. Understanding the extent of co-circulation of NA- and EU-genotype viruses across North America is important to quantify opportunities for coinfection, and, consequently, new reassortment events.

### 4.3 Mixed infections of ISAVs

The identification of mixed ISAV infections is epidemiologically important because reassortment requires different viruses to co-infect the same cell (Lowen, 2018). Detection of two lineages in the same tissue does not prove cellular coinfection, as studies of influenza A virus (a related orthomyxovirus) have shown that superinfection exclusion can spatially segregate viral populations within tissues and limit coinfection at the cellular level (Sims et al., 2023), yet establishes an opportunity for cellular coinfection and hence reassortment.

Mixed ISAV infections have previously been reported, including the concurrent detection of HPR0 and HPRdel variants within individual fish (Cárdenas et al., 2014; Romero et al., 2022; Rounsville et al., 2024) and the co-occurrence of genetically distinct HPR0 lineages within individual fish (Christiansen et al., 2021). In addition, EU- and NA-genotype ISAVs have been reported to circulate simultaneously at the farm level in Atlantic Canada (Romero et al., 2022). To our knowledge, the co-occurrence here of viruses with EU- and NA-genotype parentage represents the first documented observation of such mixed infection in ISAV.

Intergenotypic reassortments have never previously been reported for ISAV. There are at least three possible explanations for this. First, the widespread use of Sanger sequencing to genotype viruses on the basis of one segment may have prevented the identification of reassortments. Second, although NA- and EU-genotype viruses co-circulate in Atlantic Canada (Romero et al., 2022), documented mixed infections involving both genotypes are uncommon, potentially limiting opportunities for intergenotypic reassortment. Third, reassortment between genotypes may generate viruses with poor fitness that fail to spread efficiently. If the reassortant virus were consistently detected in coinfection with an EU-genotype virus, this could suggest that the EU-genotype virus was acting as a “helper virus”, potentially enabling the replication or transmission of a reassortant that would otherwise have reduced fitness. However, this pattern was not observed: only 2 of the 24 reassortant viruses were detected in samples that also contained an EU-genotype virus. The low frequency of such coinfections and the persistence of the reassortant for at least two years therefore suggests that the reassortant is able to spread independently and efficiently, though we cannot rule out that transmission could be facilitated by co-infection.

Segment-specific consensus sequences cannot directly establish which eight genome segments were packaged together within individual virions in mixed samples. We therefore assigned the segment variants in the two co-infected samples to the reassortant and co-infecting viruses based on the inference that the reassortant would retain the same genome constellation observed in the 22 non-co-infected samples. This reassortant constellation was highly conserved across the dataset, with the same combination of segments recovered from viruses sampled across nine farms and two sampling years, providing strong support for this assignment. All five EU-genotype viruses detected here, including CA26-1 and CA35-1 that were detected alongside the reassortant viruses, are dissimilar in all segments to the reassortant virus. They are therefore not closely related to the parental virus of the reassortant.

### 4.4 Implications for ISAV surveillance

Identification of an EU-NA reassortant lineage demonstrates the limitations of assigning ISAV genotype from a single genome segment. Analysis of segment 3 alone would classify these viruses as EU genotype, whereas analysis of the other seven segments places them within the NA genotype. Diagnostic assays or sequencing approaches targeting single segments could therefore produce discordant genotype assignments without revealing reassortment, suggesting that reassortant ISAV lineages are likely under-recognised by single-segment surveillance (Markussen et al., 2008; Spilsberg et al., 2024). Conventional HPR sequencing remains valuable for characterising segment 6 deletion patterns and distinguishing HPR0 from HPRdel variants, but we stress that it cannot resolve the ancestry of the other genome segments or identify discordant segment histories (McBeath et al., 2009; Gagné and LeBlanc, 2018).

Target-enrichment whole-genome sequencing addresses this limitation by enabling recovery of genome-wide sequence data from both EU- and NA-genotype viruses. In this study, it allowed simultaneous identification of the reassortant genome constellation, distinct EU-genotype lineages and mixed infections, together with characterisation of the HE HPR and F protein. Such resolution is particularly valuable in Atlantic Canada, USA and Chile, where both major ISAV genotypes co-circulate (Godoy et al., 2008; Gustafson et al., 2008; Gagné and LeBlanc, 2018; Romero et al., 2022). Whole-genome sequencing could therefore complement routine diagnostics and targeted surveillance, particularly when discordant genotype assignments are obtained or mixed infections are suspected.

Although the samples analysed here were not collected through systematic regional surveillance, broader whole-genome sequencing of ISAV across the Bay of Fundy and elsewhere in Atlantic Canada will be important to determine the prevalence, geographic distribution and persistence of the reassortant lineage, including whether it continues to circulate. This is particularly important given the marked increase in disease-associated ISA cases in New Brunswick in 2021 (Canadian Food Inspection Agency, 2026), although the available data do not establish any association between this increase and the reassortant lineage identified here during the same period. Importantly, the phenotypic consequences of this reassortment remain unknown. Whether the combination of EU- and NA-genotype segments alters viral fitness, transmissibility, persistence or pathogenicity relative to non-reassortant ISAV will require experimental investigation. Expanded sequencing of both contemporary and archived ISAV-positive samples could help establish how long this lineage has been present and identify additional reassortant viruses that may have gone unrecognised by single-segment surveillance. More extensive genome-based surveillance, combined with phenotypic characterisation of reassortant viruses, would provide a more complete understanding of how genotype co-circulation and reassortment shape ISAV evolution, transmission and disease dynamics in Atlantic Canadian salmon aquaculture.

## Acknowledgements

This work was supported by a UK Research and Innovation (UKRI) Biotechnology and Biological Sciences Research Council (BBSRC) New Investigator Grant awarded to S.C.H. (BB/W006294/1). J.O.B. was supported by an Animal Care Trust Studentship from the Royal Veterinary College. The authors thank all producers who participated in the study and provided samples and associated information.

## Data availability statement

The nucleotide sequences generated in this study have been deposited in the NCBI GenBank database under accession numbers QB278664 - QB278906.

## References

Aldrin, M., Storvik, B., Frigessi, A., Viljugrein, H. and Jansen, P.A. (2010). A stochastic model for the assessment of the transmission pathways of heart and skeleton muscle inflammation, pancreas disease and infectious salmon anaemia in marine fish farms in Norway. Preventive Veterinary Medicine, 101, 51–61. doi: 10.1016/j.prevetmed.2009.09.010.

Bates, D.M. and Watts, D.G. (1988). Nonlinear Regression Analysis and Its Applications. Wiley, New York. doi: 10.1002/9780470316757.

Canadian Food Inspection Agency (2026). Locations infected with infectious salmon anaemia. Available at: https://inspection.canada.ca/en/animal-health/aquatic-animals/diseases/reportable-diseases/isa/locations-infected (accessed 20 July 2026).

Cárdenas, C., Carmona, M., Gallardo, A., Labra, A. and Marshall, S.H. (2014). Coexistence in field samples of two variants of the infectious salmon anemia virus: a putative shift to pathogenicity. PLoS ONE, 9, e87832. doi: 10.1371/journal.pone.0087832.

Cárdenas, M., Michelson, S., Pérez, D.R., Montoya, M., Toledo, J., Vásquez-Martínez, Y. and Cortez-San Martín, M. (2022). Infectious salmon anemia virus infectivity is determined by multiple segments with an important contribution from segment 5. Viruses, 14, 631. doi: 10.3390/v14030631.

Castro-Nallar, E., Pérez-Losada, M., Burton, G.F. and Crandall, K.A. (2011). Molecular phylodynamics and protein modeling of infectious salmon anemia virus (ISAV). BMC Evolutionary Biology, 11, 349. doi: 10.1186/1471-2148-11-349.

Christiansen, D.H., McBeath, A.J.A., Aamelfot, M., Matejusova, I., Fourrier, M., White, P., Petersen, P.E. and Falk, K. (2017). First field evidence of the evolution from a non-virulent HPR0 to a virulent HPRdel infectious salmon anaemia virus. Journal of General Virology, 98, 595–606. doi: 10.1099/jgv.0.000741.

Christiansen, D.H., Petersen, P.E., Dahl, M.M., Vest, N., Aamelfot, M., Kristoffersen, A.B., Jansen, M.D., Matejusova, I., Gallagher, M.D., Jónsson, G., Rodriguez, E., Fosse, J.H. and Falk, K. (2021). No evidence of the vertical transmission of non-virulent infectious salmon anaemia virus (ISAV-HPR0) in farmed Atlantic salmon. Viruses, 13, 2428. doi: 10.3390/v13122428.

Cottet, L., Rivas-Aravena, A., Cortez-San Martín, M., Sandino, A.M. and Spencer, E. (2010). Bioinformatic analysis of the genome of infectious salmon anemia virus associated with outbreaks with high mortality in Chile. Journal of Virology, 84, 11916–11928. doi: 10.1128/JVI.01202-10.

Danecek, P., Bonfield, J.K., Liddle, J., Marshall, J., Ohan, V., Pollard, M.O., et al. (2021). Twelve years of SAMtools and BCFtools. GigaScience, 10, giab008. doi: 10.1093/gigascience/giab008.

Depledge, D.P., Palser, A.L., Watson, S.J., Lai, I.Y.C., Gray, E.R., Grant, P., et al. (2011). Specific capture and whole-genome sequencing of viruses from clinical samples. PLoS ONE, 6, e27805. doi: 10.1371/journal.pone.0027805.

Devold, M., Karlsen, M. and Nylund, A. (2006). Sequence analysis of the fusion protein gene from infectious salmon anemia virus isolates: evidence of recombination and reassortment. Journal of General Virology, 87, 2031–2040. doi: 10.1099/vir.0.81687-0.

Drummond, A.J., Ho, S.Y.W., Phillips, M.J. and Rambaut, A. (2006). Relaxed phylogenetics and dating with confidence. PLoS Biology, 4, e88. doi: 10.1371/journal.pbio.0040088.

Drummond, A.J. and Rambaut, A. (2007). BEAST: Bayesian evolutionary analysis by sampling trees. BMC Evolutionary Biology, 7, 214. doi: 10.1186/1471-2148-7-214.

Duault, H., Bailly, M., Zhao, M., Hill, S.C. and Fournié, G. (2025). Network of ship movements and its potential in spreading pathogenic infectious salmon anemia virus among Norwegian fish farms. Preventive Veterinary Medicine, 245, 106667. doi: 10.1016/j.prevetmed.2025.106667.

Edgar, R.C. (2022). Muscle5: High-accuracy alignment ensembles enable unbiased assessments of sequence homology and phylogeny. Nature Communications, 13, 6968. doi: 10.1038/s41467-022-34630-w.

Falk, K., Namork, E., Rimstad, E., Mjaaland, S. and Dannevig, B.H. (1997). Characterization of infectious salmon anemia virus, an orthomyxo-like virus isolated from Atlantic salmon (*Salmo salar* L.). Journal of Virology, 71, 9016–9023. doi: 10.1128/JVI.71.12.9016-9023.1997.

Fan, Y., Wu, R., Chen, M.-H., Kuo, L. and Lewis, P.O. (2011). Choosing among partition models in Bayesian phylogenetics. Molecular Biology and Evolution, 28, 523–532. doi: 10.1093/molbev/msq224.

Fourrier, M., Lester, K., Thoen, E., Mikalsen, A., Evensen, Ø., Falk, K., Collet, B. and McBeath, A. (2014). Deletions in the highly polymorphic region of infectious salmon anaemia virus HPR0 haemagglutinin– esterase enhance viral fusion and influence the interaction with the fusion protein. Journal of General Virology, 95, 1015–1024. doi: 10.1099/vir.0.061648-0.

Gagné, N. and LeBlanc, F. (2018). Overview of infectious salmon anaemia virus in Atlantic Canada and first report of an ISAV North American-HPR0 subtype. Journal of Fish Diseases, 41, 421–430. doi: 10.1111/jfd.12670.

Gautam, R., Price, D., Revie, C.W., Gardner, I.A., Vanderstichel, R., Gustafson, L., Klotins, K. and Beattie, M. (2018). Connectivity-based risk ranking of infectious salmon anaemia virus (ISAv) outbreaks for targeted surveillance planning in Canada and the USA. Preventive Veterinary Medicine, 159, 92–98. doi: 10.1016/j.prevetmed.2018.09.001.

Gautam, M., Hammell, K.L., Burnley, H., O’Brien, N., Whelan, D. and Thakur, K.K. (2023). Description of spatiotemporal patterns of infectious salmon anemia virus (ISAV) detections in marine Atlantic salmon farms in Newfoundland and Labrador. Journal of Aquatic Animal Health, 35, 296–307. doi: 10.1002/aah.10205.

Gill, M.S., Lemey, P., Faria, N.R., Rambaut, A., Shapiro, B. and Suchard, M.A. (2013). Improving Bayesian population dynamics inference: a coalescent-based model for multiple loci. Molecular Biology and Evolution, 30, 713–724. doi: 10.1093/molbev/mss265.

Godoy, M.G., Aedo, A., Kibenge, M.J.T., Groman, D.B., Yason, C.V., Grothusen, H., Lisperguer, A., Calbucura, M., Avendaño, F., Imilán, M., Jarpa, M. and Kibenge, F.S.B. (2008). First detection, isolation and molecular characterization of infectious salmon anaemia virus associated with clinical disease in farmed Atlantic salmon (*Salmo salar*) in Chile. BMC Veterinary Research, 4, 28. doi: 10.1186/1746-6148-4-28.

Goić, B., Bustamante, J., Miquel, A., Alvarez, M., Vera, M.I., Valenzuela, P.D.T. and Burzio, L.O. (2008). The nucleoprotein and the viral RNA of infectious salmon anemia virus are localized in the nucleolus of infected cells. Virology, 379, 55–63. doi: 10.1016/j.virol.2008.05.036.

Griffiths, R.C. and Tavaré, S. (1994). Simulating probability distributions in the coalescent. Theoretical Population Biology, 46, 131–159. doi: 10.1006/tpbi.1994.1023.

Gustafson, L., Ellis, S., Bouchard, D., Robinson, T., Marenghi, F., Warg, J. and Giray, C. (2008). Estimating diagnostic test accuracy for infectious salmon anaemia virus in Maine, USA. Journal of Fish Diseases, 31, 117–125. doi: 10.1111/j.1365-2761.2007.00873.x.

Hao, X., Hu, J., Wang, J., Xu, J., Cheng, H., Xu, Y., Li, Q., He, D., Liu, X., Wang, X., Gu, M., Hu, S., Xu, X., Liu, H., Chen, S., Peng, D. and Liu, X. (2016). Reassortant H5N1 avian influenza viruses containing PA or NP gene from an H9N2 virus significantly increase the pathogenicity in mice. Veterinary Microbiology, 192, 95–101. doi: 10.1016/j.vetmic.2016.07.002.

Hoang, D.T., Chernomor, O., von Haeseler, A., Minh, B.Q. and Vinh, L.S. (2018). UFBoot2: Improving the ultrafast bootstrap approximation. Molecular Biology and Evolution, 35, 518–522. doi: 10.1093/molbev/msx281.

Ince, W.L., Gueye-Mbaye, A., Bennink, J.R. and Yewdell, J.W. (2013). Reassortment complements spontaneous mutation in influenza A virus NP and M1 genes to accelerate adaptation to a new host. Journal of Virology, 87, 4330–4338. doi: 10.1128/JVI.02749-12.

Johnson, A., Binette, S.L., Cook-Versloot, M., Beattie, M., McGeachy, S., Gagné, N., McDonald, J.T. and Ritchie, R.J. (2008). Association between ISAV mortalities and ISAV molecular type in the Bay of Fundy, Canada. Canadian Technical Report of Fisheries and Aquatic Sciences, 2782, iv + 15 pp.

Kalyaanamoorthy, S., Minh, B.Q., Wong, T.K.F., von Haeseler, A. and Jermiin, L.S. (2017). ModelFinder: fast model selection for accurate phylogenetic estimates. Nature Methods, 14, 587–589. doi: 10.1038/nmeth.4285.

Kearse, M., Moir, R., Wilson, A., Stones-Havas, S., Cheung, M., Sturrock, S., et al. (2012). Geneious Basic: an integrated and extendable desktop software platform for the organization and analysis of sequence data. Bioinformatics, 28, 1647–1649. doi: 10.1093/bioinformatics/bts199.

Kibenge, F.S.B., Kibenge, M.J.T., Wang, Y., Qian, B., Hariharan, S. and McGeachy, S. (2007a). Mapping of putative virulence motifs on infectious salmon anemia virus surface glycoprotein genes. Journal of General Virology, 88, 3100–3111. doi: 10.1099/vir.0.83097-0.

Kibenge, F.S.B., Xu, H., Kibenge, M.J.T., Qian, B. and Joseph, T. (2007b). Characterization of gene expression on genomic segment 7 of infectious salmon anaemia virus. Virology Journal, 4, 34. doi: 10.1186/1743-422X-4-34.

Kibenge, F.S.B., Godoy, M.G., Wang, Y., Kibenge, M.J.T., Gherardelli, V., Mansilla, S., et al. (2009). Infectious salmon anaemia virus (ISAV) isolated from the ISA disease outbreaks in Chile diverged from ISAV isolates from Norway around 1996 and was disseminated around 2005, based on surface glycoprotein gene sequences. Virology Journal, 6, 88. doi: 10.1186/1743-422X-6-88.

Kibenge, M.J.T., Iwamoto, T., Wang, Y., Morton, A., Godoy, M.G. and Kibenge, F.S.B. (2016). Discovery of variant infectious salmon anaemia virus (ISAV) of European genotype in British Columbia, Canada. Virology Journal, 13, 3. doi: 10.1186/s12985-015-0459-1.

Kingman, J.F.C. (1982). On the genealogy of large populations. Journal of Applied Probability, 19A, 27–43. doi: 10.2307/3213548.

Krossøy, B., Hordvik, I., Nilsen, F., Nylund, A. and Endresen, C. (1999). The putative polymerase sequence of infectious salmon anemia virus suggests a new genus within the *Orthomyxoviridae*. Journal of Virology, 73, 2136–2142. doi: 10.1128/JVI.73.3.2136-2142.1999.

Kulshreshtha, V., Kibenge, M., Salonius, K., Simard, N., Riveroll, A. and Kibenge, F.S.B. (2010). Identification of the 3′ and 5′ terminal sequences of the 8 RNA genome segments of European and North American genotypes of infectious salmon anemia virus (an orthomyxovirus) and evidence for quasispecies based on the non-coding sequences of transcripts. Virology Journal, 7, 338. doi: 10.1186/1743-422X-7-338.

Lartillot, N. and Philippe, H. (2006). Computing Bayes factors using thermodynamic integration. Systematic Biology, 55, 195–207. doi: 10.1080/10635150500433722.

Li, H. (2018). Minimap2: pairwise alignment for nucleotide sequences. Bioinformatics, 34, 3094–3100. doi: 10.1093/bioinformatics/bty191.

Lowen, A.C. (2018). It’s in the mix: reassortment of segmented viral genomes. PLoS Pathogens, 14, e1007200. doi: 10.1371/journal.ppat.1007200.

Madhun, A.S., Mæhle, S., Wennevik, V. and Karlsbakk, E. (2019). Prevalence and genotypes of infectious salmon anaemia virus (ISAV) in returning wild Atlantic salmon (*Salmo salar* L.) in northern Norway. Journal of Fish Diseases, 42, 1217–1221. doi: 10.1111/jfd.13021.

Mardones, F.O., Martinez-Lopez, B., Valdes-Donoso, P., Carpenter, T.E. and Perez, A.M. (2014). The role of fish movements and the spread of infectious salmon anemia virus in Chile, 2007–2009. Preventive Veterinary Medicine, 114, 37–46. doi: 10.1016/j.prevetmed.2014.01.012.

Markussen, T., Jonassen, C.M., Numanovic, S., Braaen, S., Hjortaas, M., Nilsen, H. and Mjaaland, S. (2008). Evolutionary mechanisms involved in the virulence of infectious salmon anaemia virus (ISAV), a piscine orthomyxovirus. Virology, 374, 515–527. doi: 10.1016/j.virol.2008.01.019.

Martin, D.P., Varsani, A., Roumagnac, P., Botha, G., Maslamoney, S., Schwab, T., Kelz, Z., Kumar, V. and Murrell, B. (2021). RDP5: a computer program for analyzing recombination in, and removing signals of recombination from, nucleotide sequence datasets. Virus Evolution, 7, veaa087. doi: 10.1093/ve/veaa087.

McBeath, A.J.A., Bain, N. and Snow, M. (2009). Surveillance for infectious salmon anaemia virus HPR0 in marine Atlantic salmon farms across Scotland. Diseases of Aquatic Organisms, 87, 161–169. doi: 10.3354/dao02128.

McDonald, S.M., Nelson, M.I., Turner, P.E. and Patton, J.T. (2016). Reassortment in segmented RNA viruses: mechanisms and outcomes. Nature Reviews Microbiology, 14, 448–460. doi: 10.1038/nrmicro.2016.46.

Minh, B.Q., Schmidt, H.A., Chernomor, O., Schrempf, D., Woodhams, M.D., von Haeseler, A. and Lanfear, R. (2020). IQ-TREE 2: new models and efficient methods for phylogenetic inference in the genomic era. Molecular Biology and Evolution, 37, 1530–1534. doi: 10.1093/molbev/msaa015.

Mjaaland, S., Rimstad, E., Falk, K. and Dannevig, B.H. (1997). Genomic characterization of the virus causing infectious salmon anemia in Atlantic salmon (*Salmo salar* L.): an orthomyxo-like virus in a teleost. Journal of Virology, 71, 7681–7686. doi: 10.1128/JVI.71.10.7681-7686.1997.

Mjaaland, S., Hungnes, O., Teig, A., Dannevig, B.H., Thorud, K. and Rimstad, E. (2002). Polymorphism in the infectious salmon anemia virus hemagglutinin gene: importance and possible implications for evolution and ecology of infectious salmon anemia disease. Virology, 304, 379–391. doi: 10.1006/viro.2002.1658.

Mullins, J.E., Groman, D.B. and Wadowska, D. (1998). Infectious salmon anaemia in salt water Atlantic salmon (*Salmo salar* L.) in New Brunswick, Canada. Bulletin of the European Association of Fish Pathologists, 18, 110–114.

Munir, K. and Kibenge, F.S.B. (2004). Detection of infectious salmon anaemia virus by real-time RT-PCR. Journal of Virological Methods, 117, 37–47. doi: 10.1016/j.jviromet.2003.11.020.

Oxford Nanopore Technologies (2023). Medaka: sequence correction provided by ONT Research, version 1.11.3 [computer software]. GitHub. Available at: https://github.com/nanoporetech/medaka/releases/tag/v1.11.3 (accessed 6 August 2026).

Plarre, H., Devold, M., Snow, M. and Nylund, A. (2005). Prevalence of infectious salmon anaemia virus (ISAV) in wild salmonids in western Norway. Diseases of Aquatic Organisms, 66, 71–79. doi: 10.3354/dao066071.

Plarre, H., Nylund, A., Karlsen, M., Brevik, Ø., Sæther, P.A. and Vike, S. (2012). Evolution of infectious salmon anaemia virus (ISA virus). Archives of Virology, 157, 2309–2326. doi: 10.1007/s00705-012-1438-0.

Rambaut, A., Lam, T.T., Max Carvalho, L. and Pybus, O.G. (2016). Exploring the temporal structure of heterochronous sequences using TempEst (formerly Path-O-Gen). Virus Evolution, 2, vew007. doi: 10.1093/ve/vew007.

Rambaut, A. (2018). FigTree v1.4.4. Institute of Evolutionary Biology, University of Edinburgh, Edinburgh, UK. Available at: https://github.com/rambaut/figtree.

Rambaut, A., Drummond, A.J., Xie, D., Baele, G. and Suchard, M.A. (2018). Posterior summarization in Bayesian phylogenetics using Tracer 1.7. Systematic Biology, 67, 901–904. doi: 10.1093/sysbio/syy032.

Rimstad, E. and Markussen, T. (2020). Infectious salmon anaemia virus—molecular biology and pathogenesis of the infection. Journal of Applied Microbiology, 129, 85–97. doi: 10.1111/jam.14567.

Romero, J.F., Gardner, I.A., Hammell, K.L., Groman, D., Whelan, D., O’Brien, N., Hawkins, L.J., Burnley, H. and Thakur, K.K. (2022). Descriptive epidemiology of variants of infectious salmon anaemia virus in four Atlantic salmon farms in Newfoundland and Labrador, Canada. Journal of Fish Diseases, 45, 919–930. doi: 10.1111/jfd.13617.

Rounsville, T.F., Polinski, M.P., Marini, A.G., Turner, S.M., Vendramin, N., Cuenca, A., Pietrak, M., Peterson, B.C. and Bouchard, D.A. (2024). Rapid differentiation of infectious salmon anemia virus avirulent (HPR0) from virulent (HPRΔ) variants using multiplex RT-qPCR. Journal of Veterinary Diagnostic Investigation, 36, 329–337. doi: 10.1177/10406387231223290.

Sims, A., Tornaletti, L.B., Jasim, S., Pirillo, C., Devlin, R., Hirst, J.C., Loney, C., Wojtus, J., Sloan, E., Thorley, L., Boutell, C., Roberts, E. and Hutchinson, E. (2023). Superinfection exclusion creates spatially distinct influenza virus populations. PLoS Biology, 21, e3001941. doi: 10.1371/journal.pbio.3001941.

Spearman, C. (1904). The proof and measurement of association between two things. The American Journal of Psychology, 15, 72–101. doi: 10.2307/1412159.

Spilsberg, B.S., Leithaug, M.L., Christiansen, D.H., Dahl, M.M., Petersen, P.E., Lagesen, K., Fiskebeck, E.M.L.Z., Moldal, T. and Boye, M. (2024). Development and application of a whole genome amplicon sequencing method for infectious salmon anemia virus (ISAV). Frontiers in Microbiology, 15, 1392607. doi: 10.3389/fmicb.2024.1392607.

Suchard, M.A., Lemey, P., Baele, G., Ayres, D.L., Drummond, A.J. and Rambaut, A. (2018). Bayesian phylogenetic and phylodynamic data integration using BEAST 1.10. Virus Evolution, 4, vey016. doi: 10.1093/ve/vey016.

Tavaré, S. (1986). Some probabilistic and statistical problems in the analysis of DNA sequences. In: Miura, R.M. (ed.), Some Mathematical Questions in Biology—DNA Sequence Analysis. Lectures on Mathematics in the Life Sciences, vol. 17. American Mathematical Society, Providence, pp. 57–86.

Thomas, M.N., Ciacci Zanella, G., Cowan, B., Caceres, C.J., Rajao, D.S., Perez, D.R., Gauger, P.C., Vincent Baker, A.L. and Anderson, T.K. (2024). Nucleoprotein reassortment enhanced transmissibility of H3 1990.4.a clade influenza A virus in swine. Journal of Virology, 98, e01703–23. doi: 10.1128/jvi.01703-23.

Valero, Y. and Cuesta, A. (2023). Reassortant viruses threatening fish aquaculture. Reviews in Aquaculture, 15, 1720–1731. doi: 10.1111/raq.12813.

Wasilenko, J.L., Lee, C.W., Sarmento, L., Spackman, E., Kapczynski, D.R., Suarez, D.L. and Pantin-Jackwood, M.J. (2008). NP, PB1, and PB2 viral genes contribute to altered replication of H5N1 avian influenza viruses in chickens. Journal of Virology, 82, 4544–4553. doi: 10.1128/JVI.02642-07.

Weitemier, K., Straub, S.C.K., Cronn, R.C., Fishbein, M., Schmickl, R., McDonnell, A. and Liston, A. (2014). Hyb-Seq: combining target enrichment and genome skimming for plant phylogenomics. Applications in Plant Sciences, 2, 1400042. doi: 10.3732/apps.1400042.

Wick, R.R. (2018). Porechop v0.2.4: adapter trimmer for Oxford Nanopore reads. GitHub. Available at: https://github.com/rrwick/Porechop (accessed 20 July 2026).

Wickham, H. (2016). ggplot2: Elegant Graphics for Data Analysis. Springer.

Xie, W., Lewis, P.O., Fan, Y., Kuo, L. and Chen, M.-H. (2011). Improving marginal likelihood estimation for Bayesian phylogenetic model selection. Systematic Biology, 60, 150–160. doi: 10.1093/sysbio/syq085.

Yang, Z. (1994). Maximum likelihood phylogenetic estimation from DNA sequences with variable rates over sites: approximate methods. Journal of Molecular Evolution, 39, 306–314. doi: 10.1007/BF00160154.

Yu, G., Smith, D.K., Zhu, H., Guan, Y. and Lam, T.T.-Y. (2017). ggtree: an R package for visualization and annotation of phylogenetic trees with their covariates and other associated data. Methods in Ecology and Evolution, 8, 28–36. doi: 10.1111/2041-210X.12628.

Zhao, M., Duault, H., Fournié, G., Solheim, M.A., Alexandersen, S., Matthews, C., Soares, S., Madhun, A., Karlsen, M. and Hill, S.C. (2026). Molecular epidemiology and multi-scale drivers of piscine myocarditis virus dispersal in salmon aquaculture. Virus Evolution, 12, veag020. doi: 10.1093/ve/veag020.

Zheng, W., Olson, J., Vakharia, V. and Tao, Y.J. (2013). The crystal structure and RNA-binding of an orthomyxovirus nucleoprotein. PLoS Pathogens, 9, e1003624. doi: 10.1371/journal.ppat.1003624.

